# Spatial transcriptomics identifies L6 corticothalamic and L6b subplate-like heterotopic neurons in an epileptogenic SLC35A2-associated brain malformation

**DOI:** 10.64898/2026.09.04.749069

**Authors:** Nao KN Tabe, Satoshi Miyashita, Kaoru Yagita, Keiya Iijima, Kazumi Shimaoka, Ayane Hosaka, Sota Okumiya, Kayo Nishitani, Kanako Komatsu, Emi Usukura, Kumiko Murayama, Terunori Sano, Masaki Takao, Masaki Iwasaki, Mikio Hoshino

## Abstract

Increased white matter (WM) heterotopic neurons are a pathological feature observed in various neurological and psychiatric disorders. Mild malformation of cortical development with oligodendroglial hyperplasia in epilepsy (MOGHE) is a drug-resistant epilepsy-associated cortical malformation characterized by increased WM heterotopic neurons and OLIG2-positive cells and frequently associated with somatic *SLC35A2* mutations. However, the precise cellular identity of these heterotopic neurons remains unclear. Using imaging-based spatial transcriptomics (iST) of *SLC35A2*-mutated MOGHE tissues, we found that most WM heterotopic neurons exhibited molecular signatures of excitatory neurons. Further analyses revealed that these heterotopic excitatory neurons included prominent populations with layer 6 corticothalamic (CT)-like and layer 6b subplate neuron (SpN)-like molecular features. These findings were further supported by human immunohistochemistry and a *Slc35a2* conditional knockout (cKO) mouse model. Together, our results provide a cellular characterization of WM heterotopic neurons in MOGHE and suggest a new perspective on disease pathophysiology centered on deep-layer excitatory neuronal populations.

## Main

White matter (WM) has traditionally been regarded as a conduit for long-range axonal communication. However, histopathological studies have documented the presence of neurons ectopically located within the WM, commonly referred to as heterotopic neurons (also termed interstitial white matter neurons, IWMN).^1,2^ Although these neurons are present in small numbers in the neurotypical human brain, increased numbers have been reported in a range of neurological and psychiatric disorders, including epilepsy, autism spectrum disorder, schizophrenia and Alzheimer disease.^3–6^ Nevertheless, why heterotopic neurons increase in these conditions, whether they represent a primary pathogenic driver or a secondary consequence of disease, and how they relate to disease mechanisms̶particularly epileptogenesis̶remain largely unresolved.

Mild malformation of cortical development with oligodendroglial hyperplasia in epilepsy (MOGHE) is a recently defined subtype of malformations of cortical development (MCD). Histopathologically, MOGHE is characterized by an excessive presence of heterotopic neurons within the WM, hyperplasia of OLIG2-positive cells, and blurring of the gray-white matter (GM-WM) junction due to the increased number of neurons in the adjacent WM.^7^ Approximately 45-73% of MOGHE cases have been reported to harbor somatic mutations in *SLC35A2*, a gene encoding a UDP-galactose transporter in the Golgi apparatus.^8,9^ Importantly, clinical and imaging studies have implicated lesion areas containing WM heterotopic neurons as epileptogenic foci, although the specific role of these neurons remains unclear.^10,11^ A recent study provided new insights into their molecular characteristics; however, their precise cellular identity has yet to be defined.^12^ In a mouse model involving conditional disruption of *Slc35a2* in dorsal pallium-derived cells, impaired neuronal migration and spontaneous epilepsy have been reported, although the presence and characteristics of WM heterotopic neurons were not directly examined in this model.^13^

In this study, we applied imaging-based spatial transcriptomics (iST) to surgical brain samples from patients with drug-resistant epilepsy who were histopathologically diagnosed with MOGHE and harbored *SLC35A2* mutations. By enabling high-resolution transcriptomic profiling while preserving tissue architecture, iST allowed us to directly characterize WM heterotopic neurons in MOGHE. We aimed to define their cellular identity and molecular features and to provide insight into their potential relationship to epileptogenic pathology.

## Results

### 1. Xenium spatial transcriptomic profiling

In this study, we enrolled two patients with drug-resistant epilepsy who were histopathologically diagnosed with MOGHE and two patients with mesial temporal lobe epilepsy (mTLE) associated with hippocampal sclerosis (HS). Somatic mutations in *SLC35A2* were identified in lesions surgically resected from both MOGHE patients. Of note, variants in *AKT1S1* (missense) and *MTOR* (intronic) were also identified in the respective MOGHE patients (Table 1).

**Table 1:** Patient information. Patient information for all human brain samples analyzed in this study, including histopathology, age and sex at surgery, cortical region analyzed, genetic findings, tissue type, and application to Xenium analysis and immunohistochemistry (IHC).

| Sample ID | Histopathology | Age | Sex | Cortical region analyzed | Genetic findings | Tissue type | Xenium | IHC |
| --- | --- | --- | --- | --- | --- | --- | --- | --- |
| C1 | HS | 4 | F | TL | NA | FFPE | O | O |
| C2 | HS | 11 | F | TL | NA | FFPE | O | O |
| M1 | MOGHE | 3 | F | TL | *SLC35A2* p.Leu120Hisfs*7,<br>*AKT1S1* p.Pro142Arg | FFPE | O | O |
| M2 | MOGHE | 14 | F | FL | *SLC35A2* p.Cys210Tyr,<br>*MTOR* c.5246+9CC>GT | FFPE | O | O |
| M3 | MOGHE | 3 | M | TL | *SLC35A2* p.Leu120Hisfs*7 | FFPE | – | O |
| M4 | MOGHE | 2 | M | TL | *SLC35A2* p.Ser209del | FFPE | – | O |

Formalin-fixed paraffin-embedded (FFPE) sections from the temporal or frontal cortex of MOGHE patients were analyzed together with control samples, which consisted of histopathologically normal temporal cortical regions from age- and sex-matched mTLE patients with HS corresponding to each MOGHE case. iST profiling was performed using the Xenium Prime 5K Human Pan Tissue and Pathways Panel (5,001 gene probes; 10X Genomics, USA). Following spatial transcriptomic analysis, the same tissue sections were subjected to H&E staining to visualize cortical architecture. This integrated approach enabled high-resolution characterization of both the spatial molecular landscape and histopathological features of MOGHE.

We first characterized histopathological features of MOGHE cases using H&E staining and immunostaining for NeuN and OLIG2 (Fig. 1A and 1B). In control cortices, cortical lamination was preserved, with NeuN-positive neurons largely confined to the GM, resulting in a clear GM-WM junction. In contrast, MOGHE cortices exhibited blurring of the GM-WM junction resulting from an increased number of heterotopic neurons in the WM. We also confirmed hyperplasia of OLIG2-positive cells in the WM of MOGHE cortices. These pathohistological features were consistently observed in the MOGHE cases enrolled in this study (Supplementary Fig. S1A-D; Table 1; Supplementary Table S1 and S2).

**Fig. 1:**
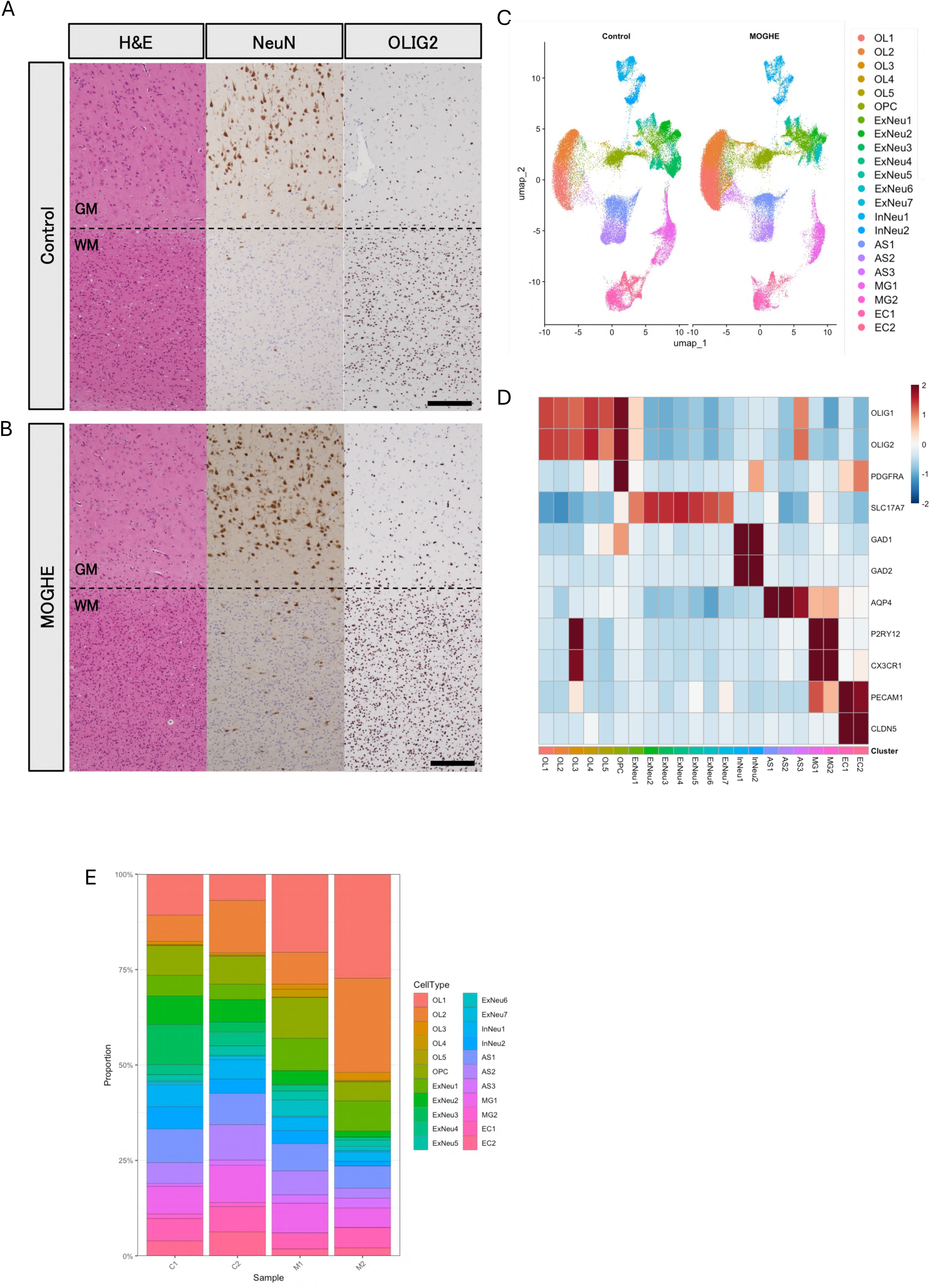
Histopathological and spatial transcriptomic overview of MOGHE and control cortices. A, B Representative images of H&E staining and immunohistochemistry for NeuN and OLIG2 in control and MOGHE cortical samples. Dashed lines indicate the gray matter (GM)-white matter (WM) junction. Scale bars, 200 µm. Images are representative of one control (C2) and one MOGHE (M1) case. C Uniform manifold approximation and projection (UMAP) visualization of major cell types identified by imaging-based spatial transcriptomic (iST) data (n = 176,800 cells). D Heatmap showing scaled expression of canonical marker genes used to annotate major cell types. E Proportions of major cell types across samples.

Our iST analysis yielded median gene counts ranging from 173 to 277 and median transcript counts ranging from 205 to 343 per cell, supporting the reliability of subsequent analyses (Supplementary Table S3). After quality-control filtering, 40,195-54,425 cells per sample were retained for downstream analyses. Unsupervised clustering following dimensionality reduction identified 22 transcriptionally distinct cell populations. Expression of canonical marker genes confirmed that iST robustly resolved major human brain cell types, including oligodendrocytes (OL1-5; *OLIG1/OLIG2*), oligodendrocyte precursor cells (OPC; *PDGFRA*), excitatory neurons (ExNeu1-7; *SLC17A7*), inhibitory neurons (InNeu1-2; *GAD1/GAD2*), astrocytes (AS1-3; *AQP4*), microglia (MG1 and MG2; *P2RY12/CX3CR1*) and endothelial cells (EC1 and EC2; *PECAM1/CLDN5*) (Fig. 1C, D; Table 2).^14–17^ At the level of broad cell-type classes, all major populations were represented across all samples, without gross shifts in overall cellular compositions that would preclude downstream comparative analyses (Fig. 1E). Together, these data established a spatially resolved cellular framework that is suitable for further investigating the histopathological and pathophysiological features of MOGHE.

**Table 2:** Cell counts for major cell types.

### 2. Molecular identity of heterotopic neurons in MOGHE revealed by spatial transcriptomics

WM heterotopic neurons are one of the hallmark pathological features of MOGHE.^7^ However, their precise cellular identity remains unclear.^12^ Given that excitatory neurons constitute the major neuronal population in the cerebral cortex, we first focused on excitatory neurons to characterize the cellular and molecular features of heterotopic neurons in MOGHE.

Subclustering of excitatory neurons (ExNeu1-ExNeu7; Fig. 1C) further identified 18 transcriptionally distinct subclusters. Of these, 13 were assigned cortical laminar identities. Four subclusters expressed glia markers and were excluded from subsequent analyses because they were considered likely to represent contaminating populations, possibly due to segmentation failure. One subcluster remained undetermined. The excitatory neuron subclusters were then annotated based on the expression of layer-associated marker genes and their spatial distribution across cortical layers. Markers used to support the annotation included *CUX2* (L2/3), *RORB* (L4), *ETV1* (L5), *THEMIS* and *OPRK1* (L6 intratelencephalic neurons; L6 IT), *FOXP2* and *TRPM3* (L6 corticothalamic neurons; L6 CT), as well as *CCN2*, *CPLX3*, and *NXPH4* (L6b subplate neurons; SpN) (Fig. 2A and 2B; Fig. S2A-C; Table 3).^14,18^ Most subclusters were shared across samples. Exceptions were L2/3 neurons, which formed sample-specific subclusters (L2/3 (C1), L2/3 (C2) and L2/3 (M1/M2)), and L6 IT-2 neurons, which formed condition-specific subclusters (L6 IT-2 (C1/C2) and L6 IT-2 (M1/M2)).

**Fig. 2:**
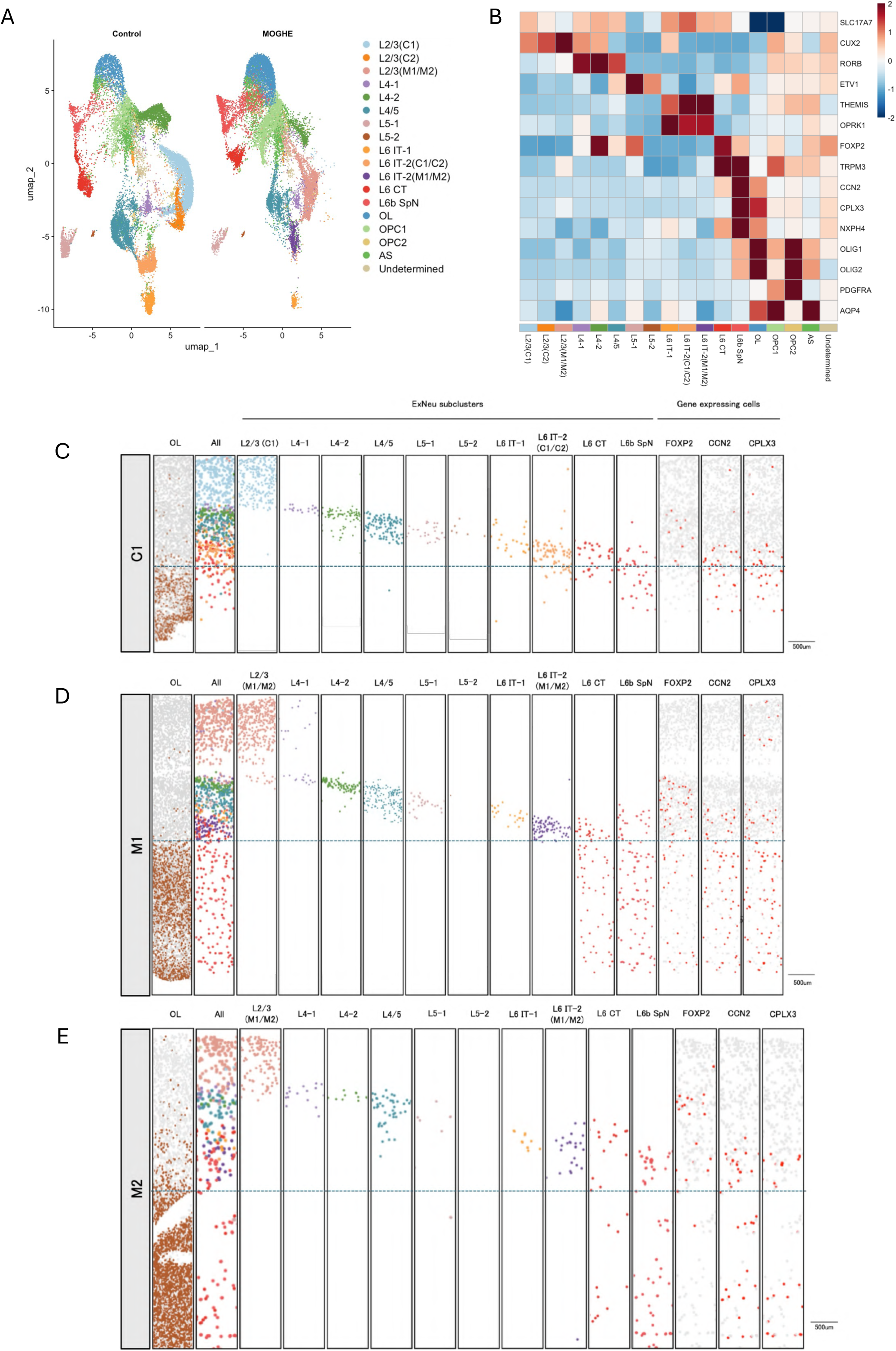
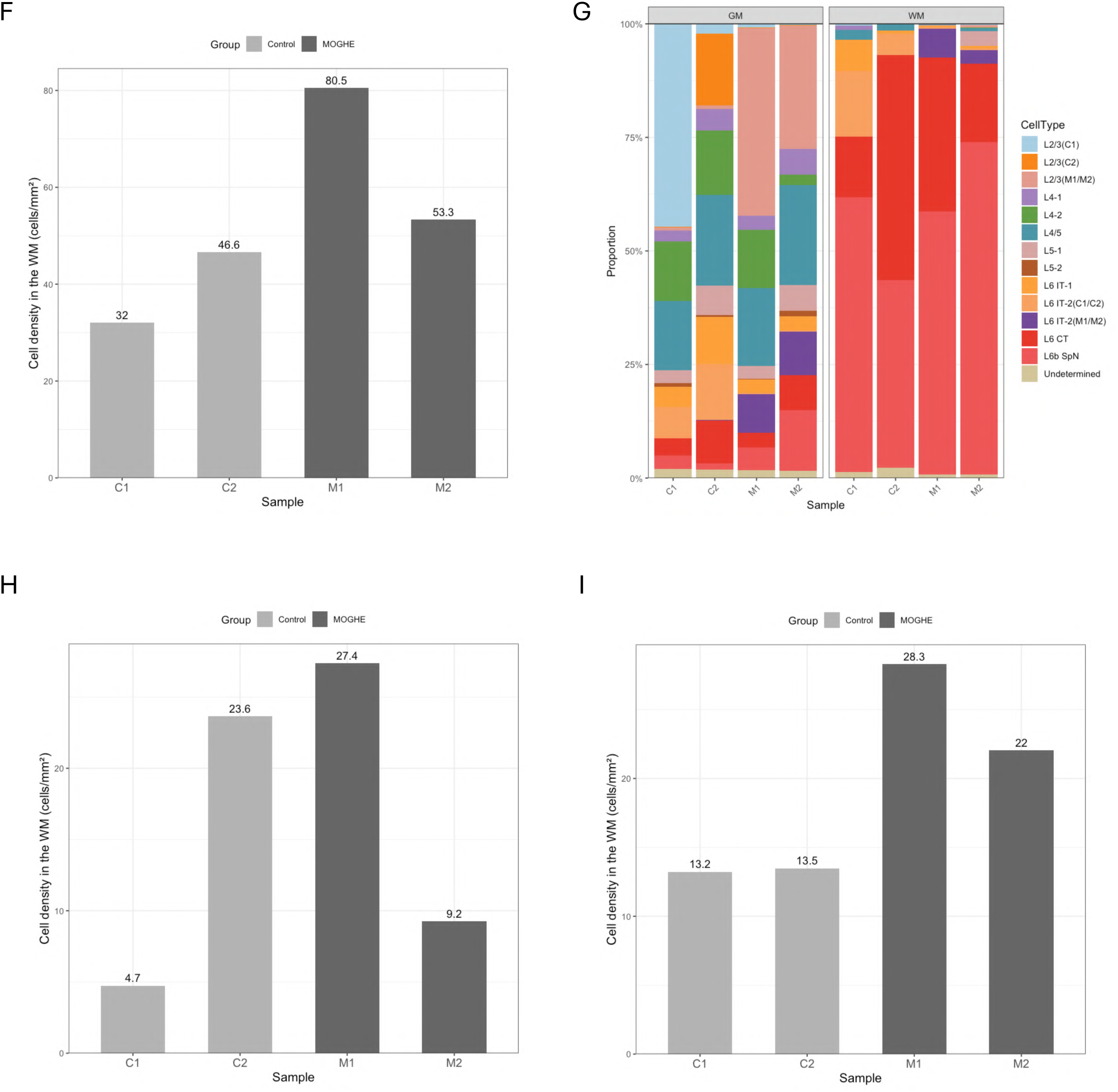
Molecular identity of heterotopic excitatory neurons in MOGHE revealed by iST analysis. A UMAP visualization of excitatory neuron subclusters identified by subclustering of the initial excitatory neuron (ExNeu) clusters (ExNeu1-7). B Heatmap showing the scaled expression of layer-associated marker genes across excitatory neuron subclusters. C-E Spatial maps of excitatory neuron subclusters in representative control (C1) and MOGHE (M1, M2) cortical sections. Dashed lines indicate the GM-WM junction. From left to right, panels show: (i) spatial distribution of oligodendrocyte (OL) clusters (OL1-OL5) identified by the initial clustering, included to delineate GM and WM compartments; (ii) spatial maps of all excitatory neuron subclusters; (iii) spatial maps of individual excitatory neuron subclusters; and (iv) spatial visualization of excitatory neurons expressing layer 6 corticothalamic (L6 CT)- and layer 6b subplate neuron (L6b SpN)-associated markers (FOXP2 and CCN2/CPLX3, respectively). In the L6 CT- and L6b SpN-associated marker expression maps, cells with marker expression above a defined threshold (> 0.5) are highlighted in red, whereas cells with lower or undetectable expression are shown in gray. Four subclusters expressing glia markers were excluded. F Density of WM-localized excitatory neurons in each sample. Four subclusters expressing glia markers were excluded from cell counts in this analysis (Table 3). G Proportions of ExNeu subclusters in GM and WM across samples. Four subclusters expressing glia markers were excluded from cell counts in this analysis (Table 3). H Density of WM-localized L6 CT-like neurons in each sample, calculated from L6 CT subclusters identified by secondary clustering of L6 CT and L6b SpN populations (Supplementary Fig. S2E and S2F; Supplementary Table S4). I Density of WM-localized L6b SpN-like neurons in each sample, calculated from L6b SpN subclusters identified by secondary clustering of L6 CT and L6b SpN populations, excluding the SpN/OL mixed subcluster (Supplementary Fig. S2E and S2F; Supplementary Table S4).

**Table 3:** Cell counts for excitatory neuron subclusters.

To assess laminar organizations of control and MOGHE cortices, we visualized spatial distributions of excitatory neurons around the GM-WM junction (Fig. 2C-E; Supplementary Fig. S2D). In control cortices, excitatory neuron subclusters exhibited a well-structured laminar organization within the GM, with excitatory neurons largely confined to the GM and only a limited number observed in the WM. In contrast, MOGHE cortices showed a marked increase in excitatory neurons localized within the WM (Fig. 2F). Analysis of subcluster composition revealed that these heterotopic excitatory neurons predominantly belonged to the L6 CT and L6b SpN subclusters, a pattern consistently observed in both control and MOGHE samples (Fig. 2G).

We next quantified the density of L6 CT and L6b SpN subclusters within the GM and WM. L6 CT neurons are deep-layer projection neurons that project to the thalamus, whereas L6b SpN neurons represent a subplate-related population involved in early cortical circuit formation and thalamocortical connectivity during development.^19,20^ The densities of L6 CT-like neurons appeared lower in the GM of MOGHE samples than in controls (Supplementary Fig. S2E-G; Supplementary Table S4), whereas no consistent pattern was observed in the WM across samples (Fig. 2H). In contrast, L6b SpN-like neurons showed higher densities in the WM of MOGHE samples than in controls (Fig. 2I; Supplementary Fig. S2H). Their densities were also higher in the GM of MOGHE samples than in controls (Supplementary Fig. S2I and S2J). Spatial mapping further demonstrated that both L6 CT and L6b SpN subclusters expressed their respective cell-type-associated markers even when localized within the WM in both control and MOGHE samples (Fig. 2C-E; Supplementary Fig. S2D).

WM neurons are known to include both excitatory and inhibitory populations.^21^ We therefore performed subclustering of inhibitory neurons (InNeu1 and InNeu2; Fig. 1C) and found that the cellular composition and spatial distributions within the WM appeared similar between control and MOGHE samples (Fig. S3A-H; Supplementary Table S5). Together, these findings suggest that the increase in WM neurons observed in MOGHE is primarily attributable to deep-layer excitatory neurons including L6 CT- and L6b SpN subclusters, with the most evident increase in L6b SpN-like populations.

### 3. Oligodendroglial alterations in MOGHE revealed by spatial transcriptomics

MOGHE is histopathologically characterized by oligodendroglial hyperplasia, defined by an increased density of OLIG2-positive cells within the white matter.^7,22^ To examine the transcriptomic features of oligodendroglial lineage cells, we first performed subclustering of OL and OPC populations (OL1-5 and OPC; Fig. 1C) identified in the initial clustering of the spatial transcriptomic data (Fig. 3A).

**Fig. 3:**
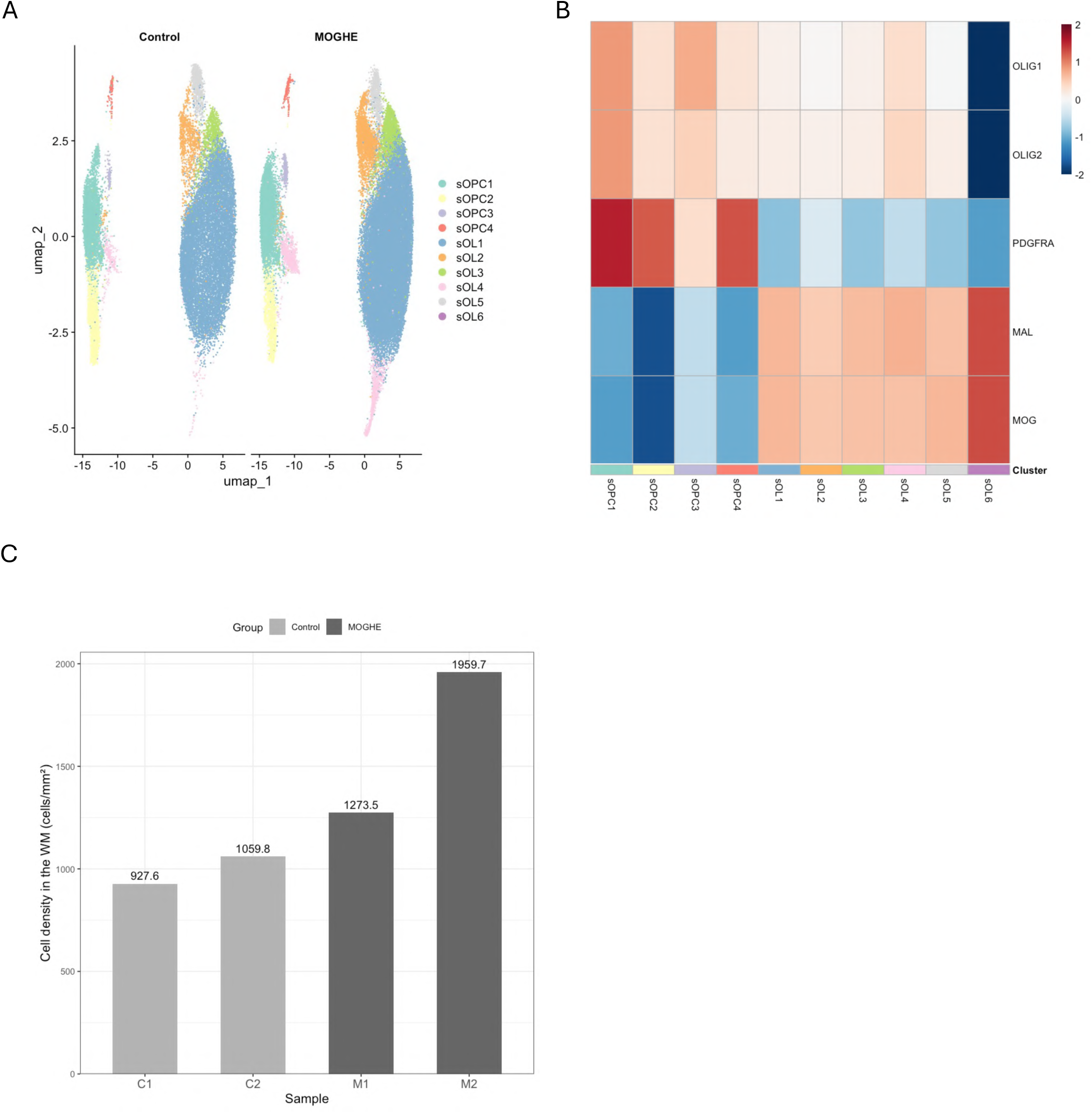
Oligodendroglial cell analysis. A UMAP visualization of OL and oligodendrocyte precursor cell (OPC) subclusters identified by subclustering of the initial OL1-OL5 and OPC clusters. B Heatmap showing the scaled expression of oligodendroglial lineage marker genes across OL and OPC subclusters. C Density of WM-localized OL and OPC populations in each sample.

OL and OPC populations were identified in both control and MOGHE samples based on the expression of canonical lineage markers. OPC subclusters (sOPC1-sOPC4) were defined by the expression of OLIG1, OLIG2, and PDGFRA, whereas oligodendrocyte subclusters (sOL1-sOL6) expressed OLIG1 and OLIG2, accompanied by more differentiated oligodendrocyte markers such as MAL and MOG (Fig. 3B).^15^ To assess whether oligodendroglial hyperplasia described histopathologically was reflected in the spatial transcriptomic data, we quantified OL and OPC densities as the number of OL and OPC-assigned cells per unit WM area. The overall density of OL and OPC populations appeared higher in MOGHE samples than in controls (Fig. 3C; Supplementary Fig. S4A and S4B), consistent with the histopathological feature of oligodendroglial hyperplasia. We did not observe apparent differences in relative subcluster composition between MOGHE and control samples (Supplementary Fig. S4C and S4D; Table 4).

**Table 4:** Cell counts for oligodendrocyte (OL) and oligodendrocyte precursor cell (OPC) subclusters.

To further characterize transcriptional alterations in oligodendroglial lineage cells associated with MOGHE, we performed differential expression analysis comparing OL and OPC populations between MOGHE and control samples. This analysis identified differentially expressed genes involved in cholesterol biosynthesis and lipid metabolism in MOGHE-associated oligodendroglial populations, including *DHCR24*, *SQLE*, *LDLR, PNPLA3* and *FABP3* (Supplementary Table S6).^23,24^ Together, these findings indicated that oligodendroglial abnormalities in MOGHE are accompanied by transcriptional alterations within oligodendroglial lineage cells.

We also performed subclustering analyses of other glial populations including astrocytes (AS1-AS3; Fig. 1C) and microglia (MG1 and MG2; Fig. 1C). Although five astrocyte subclusters and seven microglia subclusters were identified, their distributions varied across samples, with no consistent condition-associated differences apparent for most subclusters (Fig. S5A-D; Supplementary Table S7 and S8).

### 4. Immunohistochemical validation of L6 CT- and L6b SpN-like WM heterotopic neurons in human MOGHE cortex

To validate the molecular identity of heterotopic neurons revealed by spatial transcriptomic analyses, we performed immunohistochemical staining of human cortical sections from MOGHE patients using the L6 CT-associated neuron marker FOXP2 and the L6b SpN-associated markers CTGF (encoded by *CCN2*) and CPLX3.^14,25^

In control cortices, FOXP2-, CTGF- and CPLX3-positive neurons were observed in the deep cortical layers, and a subset of these marker-positive neurons was also detected within the WM. In MOGHE cortices, FOXP2-, CTGF- and CPLX3-positive neurons were likewise observed within the WM (Fig. 4A-F; Supplementary Fig. S6A-D). Similar findings were consistently observed in independent MOGHE cases (n = 2) that were not used for spatial transcriptomic analyses (Fig. 4G-H; Supplementary Fig. S6E; Supplementary Table S1 and S2).

**Fig. 4:**
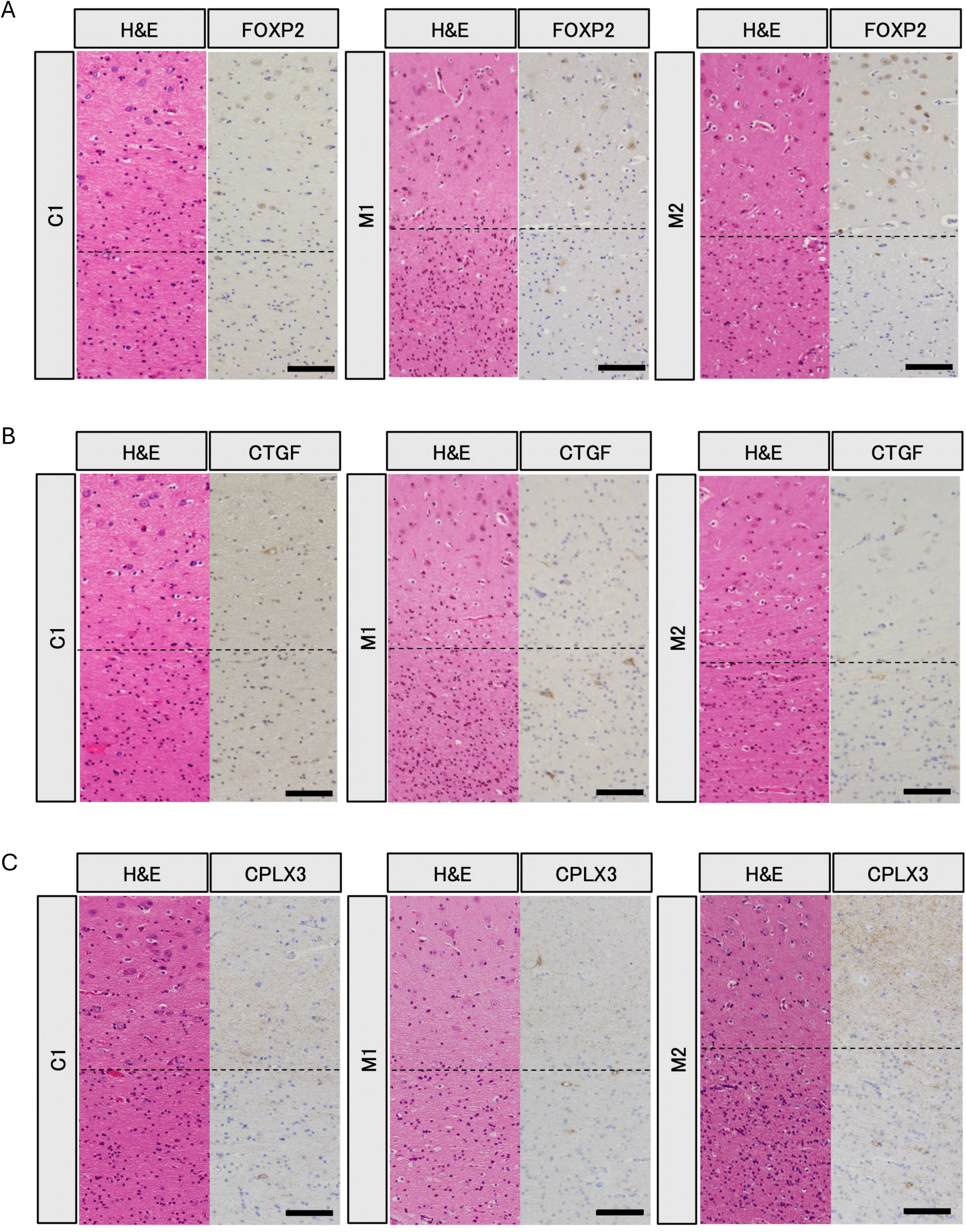
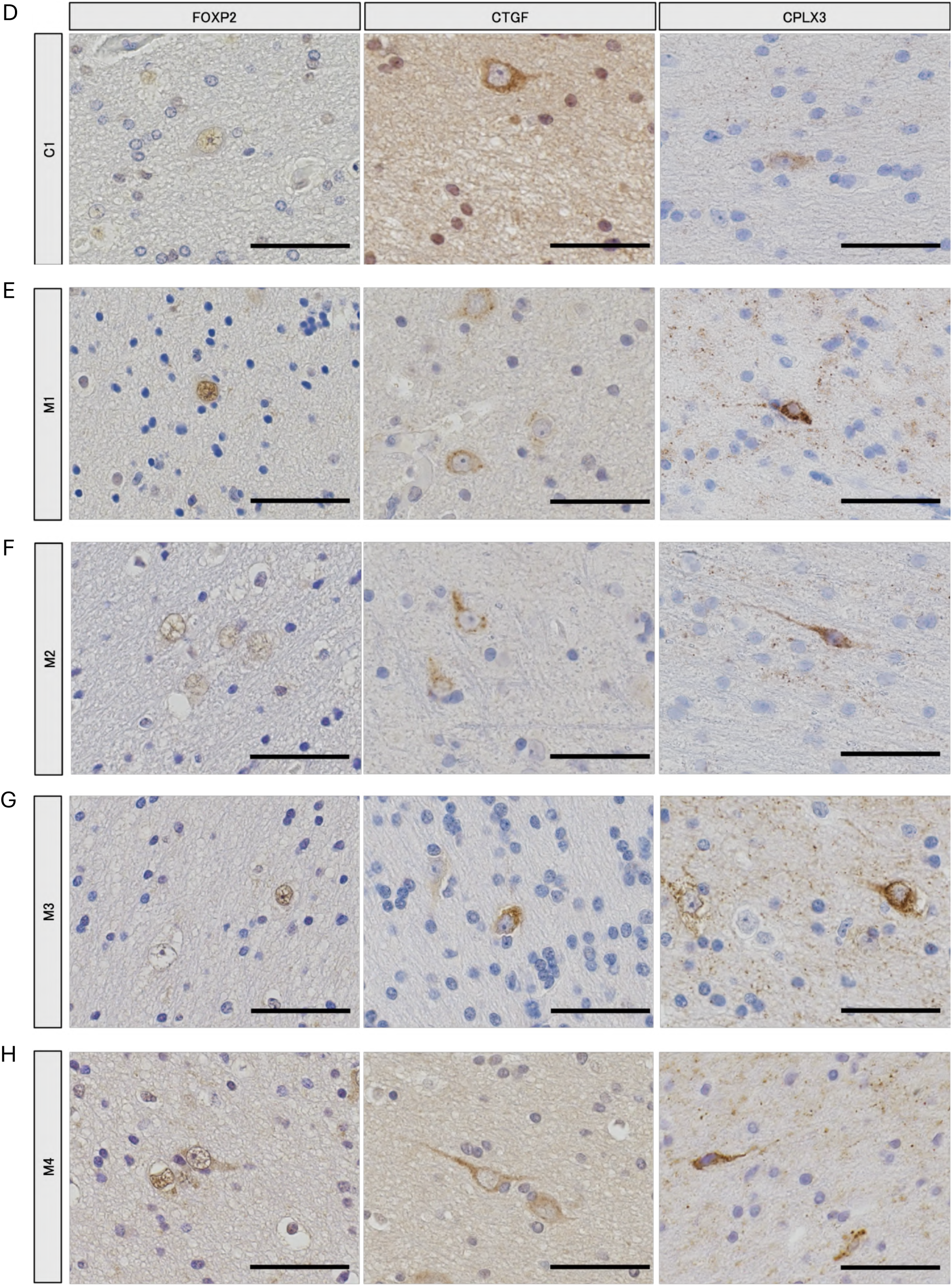
Immunohistochemical validation of L6 CT- and L6b SpN-associated markers in MOGHE. A-C Immunohistochemical staining for FOXP2 (A), CTGF (B), and CPLX3 (C) in control (C1) and MOGHE (M1, M2) cortical sections. Adjacent H&E-stained sections are shown for anatomical reference. Dashed lines indicate the GM-WM boundary. Scale bars, 100 µm. D-H Higher-magnification images of marker-positive cells in the WM regions from control (C1) and MOGHE (M1-M4) cortical sections. Scale bars, 50 µm.

The distribution patterns of FOXP2-, CTGF- and CPLX3-positive neurons were consistent with the spatial transcriptomic findings, supporting the interpretation that heterotopic neurons in MOGHE include populations with L6 CT and L6b SpN molecular identities.

### 5. Increased WM heterotopic neurons with L6 CT- and L6b SpN-like identity in a *Slc35a2* conditional knockout mouse model

To determine whether the neuronal features observed in human MOGHE cortex are recapitulated in an experimental model, we analyzed a *Slc35a2* conditional knockout (cKO) mouse generated using the Emx1-Cre driver, which targets dorsal pallium-derived excitatory neuron lineages.^26^

Quantitative assessment showed that the density of WM-localized neurons was increased in the cingulum (cg) of cKO mice, indicating an increased number of heterotopic neurons in the WM (Fig. 5A-C). To assess their molecular identity, we next performed immunohistochemical staining for markers associated with L6 CT neurons (TLE4) and L6b SpN neurons (CPLX3).^27,28^ Quantitative analysis demonstrated increased densities of both TLE4-positive and CPLX3-positive cells within the WM of cKO mice (Fig. 5D-I). Together, these findings indicate that heterotopic neurons present in the WM of *Slc35a2* cKO mice exhibit molecular features associated with L6 CT and L6b SpN populations, consistent with the molecular identities identified by human iST analysis.

**Fig. 5:**
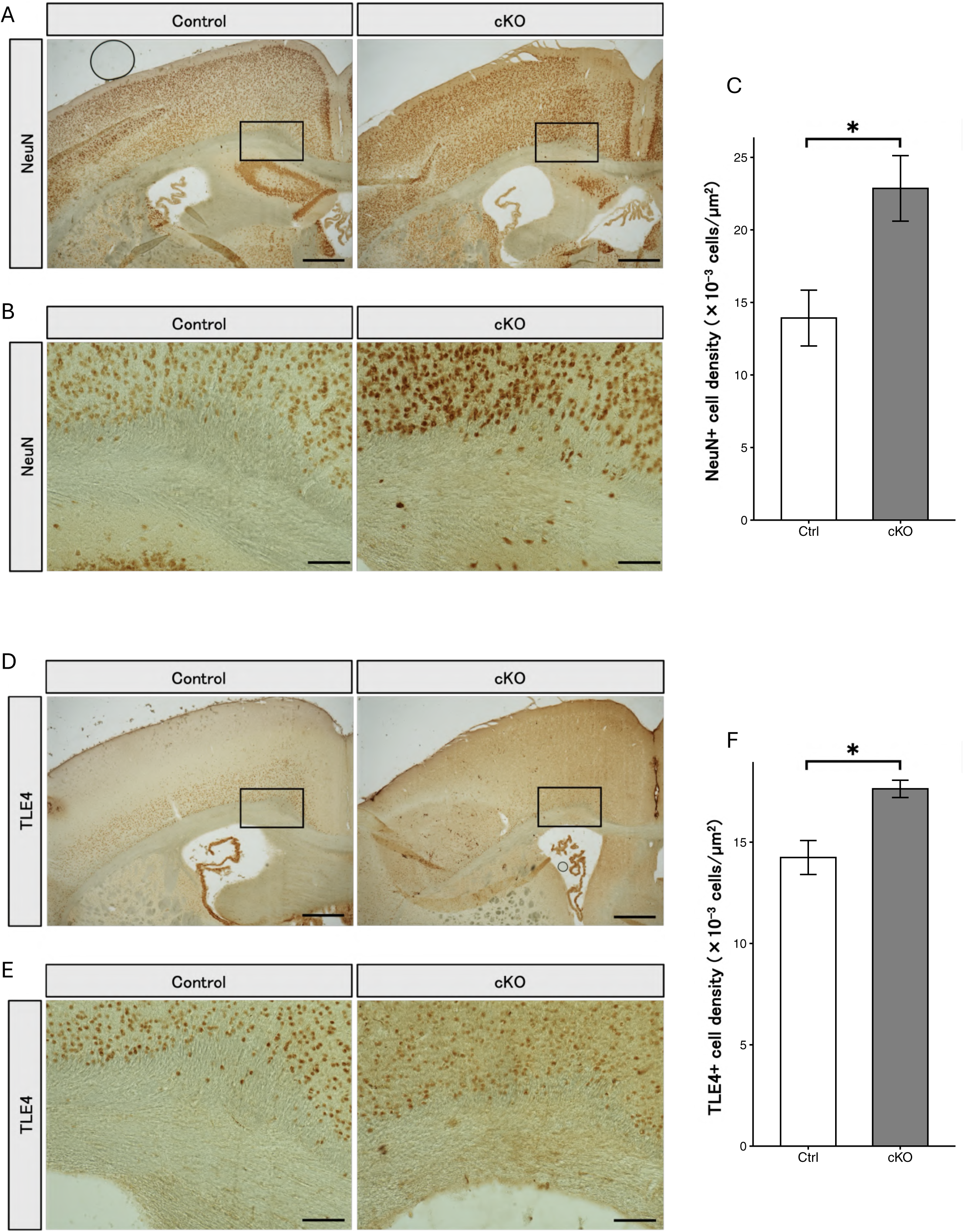
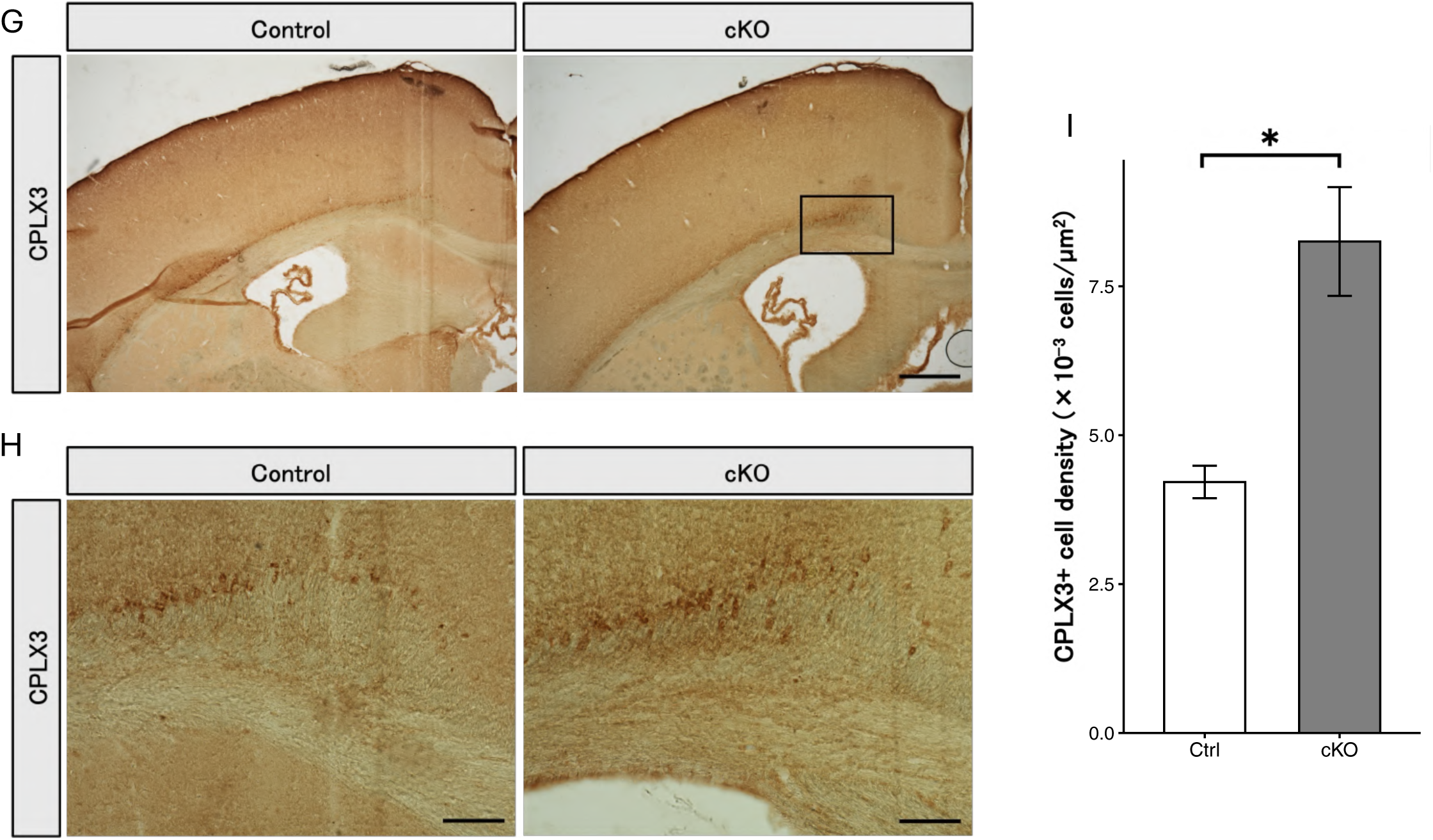
WM heterotopic neurons with L6 CT and L6b SpN-like features in a Slc35a2 conditional knockout (cKO) mouse model. A Representative NeuN immunohistochemistry in the cingulum (cg) of control and cKO mice. Scale bars, 500 µm. B Higher-magnification images of the WM regions shown in A. Scale bars, 100 µm. C Quantification of WM-localized NeuN-positive neuronal density in control and cKO mice. n = 5 control and 5 cKO mice. Data are presented as mean ± standard error of the mean (SEM). P values were calculated using Welch’s t-test. *p < 0.05. D Representative TLE4 immunohistochemistry in the cg of control and cKO mice. Scale bars, 500 µm. E Higher-magnification images of the WM regions shown in D. Scale bars, 100 µm. F Quantification of WM-localized TLE4-positive neuronal density in control and cKO mice. n = 3 control and 3 cKO mice. Data are presented as mean ± SEM. P values were calculated using Welch’s t-test. *p < 0.05. G Representative CPLX3 immunohistochemistry in the cg of control and cKO mice. Scale bars, 500 µm. H Higher-magnification images of the WM regions shown in G. Scale bars, 100 µm. I Quantification of WM-localized CPLX3-positive neuronal density in control and cKO mice. n = 3 control and 3 cKO mice. Data are presented as mean ± SEM. P values were calculated using Welch’s t-test. *p < 0.05.

## Discussion

MOGHE is a malformation of cortical development characterized by increased heterotopic neurons and hyperplasia of OLIG2-positive cells within the WM.^7^ Although WM heterotopic neurons represent a hallmark pathological feature of MOGHE, their precise cellular identity has remained unclear. In the present study, we applied iST analysis to surgically resected MOGHE tissues and demonstrated that heterotopic excitatory neurons exhibit molecular signatures of L6 CT and L6b SpN neurons. These findings were further supported by immunohistochemical analyses of human tissues and by analysis of a *Slc35a2* cKO mouse model. Together, our results provide a cellular characterization of heterotopic neurons in MOGHE.

Importantly, we observed increased numbers of TLE4-positive L6 CT-like and CPLX3-positive L6b SpN-like neurons in the WM of *Slc35a2* cKO mice, further supporting a link between *SLC35A2* deficiency and WM heterotopic neuronal abnormalities, a key pathological feature of MOGHE lesions. Because human surgical tissues are influenced by multiple factors, including genetic background, age, brain region, and seizure history, findings from human samples alone cannot fully distinguish the effects of *SLC35A2* deficiency from other patient-specific factors. In particular, *AKT1S1* and *MTOR* variants were identified in MOGHE patients used for iST analyses, respectively, and their potential contributions to the observed neuronal phenotypes cannot be completely excluded.

Nevertheless, the observation of similar L6 CT- and L6b SpN-like abnormalities in the *Slc35a2* cKO mice supports an association between *SLC35A2* and these phenotypes. Therefore, the presence of heterotopic neurons exhibiting L6 CT- and L6b SpN-like molecular features in both human MOGHE tissues and *Slc35a2* cKO mice suggests that these neuronal abnormalities may be associated with *SLC35A2* deficiency rather than solely reflecting patient-specific factors.

A previous single-nucleus RNA sequencing study of MOGHE showed that most WM heterotopic neurons were annotated as L5/6 excitatory neurons.^12^ Notably, their differentially expressed gene list for these neurons included *FOXP2, CCN2,* and *CPLX3*. These observations are consistent with our identification of L6 CT- and L6b SpN-like molecular features in heterotopic neurons in MOGHE.

Our findings also provide insight into the potential relationship between these neuronal abnormalities and epileptogenesis in MOGHE. Previous studies have yielded differing seizure phenotypes following *Slc35a2* deletion in Olig2-lineage cells, with one study reporting normal EEG activity without a clear seizure phenotype, whereas another observed spontaneous electroclinical seizures.^13,29^ In another model, *Slc35a2* disruption in dorsal pallium-derived cells using Emx1-Cre resulted in spontaneous seizures, suggesting that abnormalities in dorsal excitatory neuron lineages may contribute to epileptogenesis.^13^ In the present study, we further show that *Slc35a2* cKO mice exhibit increased WM neurons expressing molecular features associated with L6 CT- and L6b SpN-like populations.

Furthermore, clinical and imaging studies have implicated lesion areas containing WM heterotopic neurons as epileptogenic foci, although the specific contribution of these neurons to seizure generation remains unclear.^10,11^ Together, these findings raise the possibility that heterotopic L6 CT- and L6b SpN-like neurons identified in the present study may contribute to epileptogenesis and/or seizure generation.

Although the present study does not establish a causal relationship between these neuronal populations and epileptogenesis, the identification of L6 CT- and L6b SpN-like populations among heterotopic neurons may also provide clues to the potential functional consequences of WM neuronal abnormalities in MOGHE. L6 CT neurons constitute a major source of cortical projections to the thalamus, whereas SpN neurons play critical roles in the development and maturation of cortical circuits.^19,20^ Subplate-related neurons occupy a unique developmental position within the developing white matter, where they participate in early cortical circuits and the establishment and maturation of thalamocortical connectivity.^20^ Therefore, ectopic localization of these neuronal populations within the WM may affect local circuit organization and long-range connectivity. Interestingly, the higher density of L6b SpN-like neurons in the WM of MOGHE samples was not accompanied by a reduction in their GM density; rather, their density was also higher in the GM. This distribution is not readily explained by altered spatial localization alone and may instead reflect broader abnormalities in the developmental regulation of L6b SpN-like neurons.

Future studies aimed at elucidating the developmental mechanisms and functional properties of these heterotopic neurons will be important for understanding how neuronal developmental abnormalities contribute to MOGHE pathogenesis. Notably, previous experimental models of MOGHE have provided important insights into the consequences of *SLC35A2* dysfunction in cortical development and epileptogenesis, with neuronal abnormalities primarily investigated in the context of migration of upper-layer cortical neurons. Our findings extend these studies by highlighting deep-layer neurons, including L6 CT and L6b SpN neurons, as additional neuronal populations that warrant further investigation in future experimental models.^13,30–32^

One plausible molecular mechanism linking *SLC35A2* dysfunction to WM heterotopic neuronal abnormalities is impaired glycosylation-dependent regulation of neuronal development and positioning. *SLC35A2* encodes a UDP-galactose transporter in the Golgi apparatus, and its dysfunction is known to disrupt glycosylation, leading to defective glycan modification of cell-surface and secreted glycoproteins.^33^ Consistent with this, a previous study identified abnormal N-glycosylation profiles in brain tissues harboring somatic *SLC35A2* variants^33^. More recently, Liu et al. demonstrated N-glycosylation defects associated with somatic *SLC35A2* variants in MOGHE and reported reduced N-acetyllactosamine (LacNAc) levels in WM heterotopic neurons.^34^ Such glycoproteins include molecules involved in cell-cell adhesion,^34^ extracellular matrix interactions, receptor trafficking,^35^ axon guidance, and synaptic maturation,^36,37^ all of which are relevant to cortical development. Therefore, altered glycosylation could affect the proliferation, migration, terminal positioning, and/or survival of deep-layer and subplate-related excitatory neurons, thereby contributing to the accumulation of L6 CT-like and L6b SpN-like heterotopic neuronal populations in the WM. Although glycosylation was not examined in the present study, these findings raise the possibility that glycosylation defects may act upstream of the neuronal developmental and positioning abnormalities identified here. Future studies combining lineage-specific *Slc35a2* mouse models with cell-type-resolved glycomic, glycoproteomic, and spatial transcriptomic analyses will be important to determine whether L6 CT-like and L6b SpN-like heterotopic neurons are particularly vulnerable to *SLC35A2*-dependent glycosylation defects.

Beyond MOGHE, WM neurons are known to be present in the normal and diseased human brain, and their increase in number has been reported in association with temporal lobe epilepsy, autism spectrum disorder, schizophrenia and Alzheimer’s disease.^3–6^ Consistent with our findings in MOGHE patients, previous studies have shown that subsets of WM neurons in epilepsy patients express markers associated with deep-layer cortical neurons, including TBR1, CTIP2, and TLE4, leading to the hypothesis that these cells may be related to subplate neurons or deep-layer cortical neurons and may reflect, at least in part, abnormalities arising during early stages of cortical development.^3,38^

Several limitations should be acknowledged. First, the number of human samples available for spatial transcriptomic analysis was limited. Second, although our study defines the molecular identities of heterotopic neurons in MOGHE, it does not directly address their developmental origin or functional contribution to disease pathogenesis. Future studies using cell lineage-specific experimental models will be required to determine how L6 CT and L6b SpN populations contribute to cortical malformation and epileptogenesis in MOGHE.

---

**Supplementary Fig. S1:**
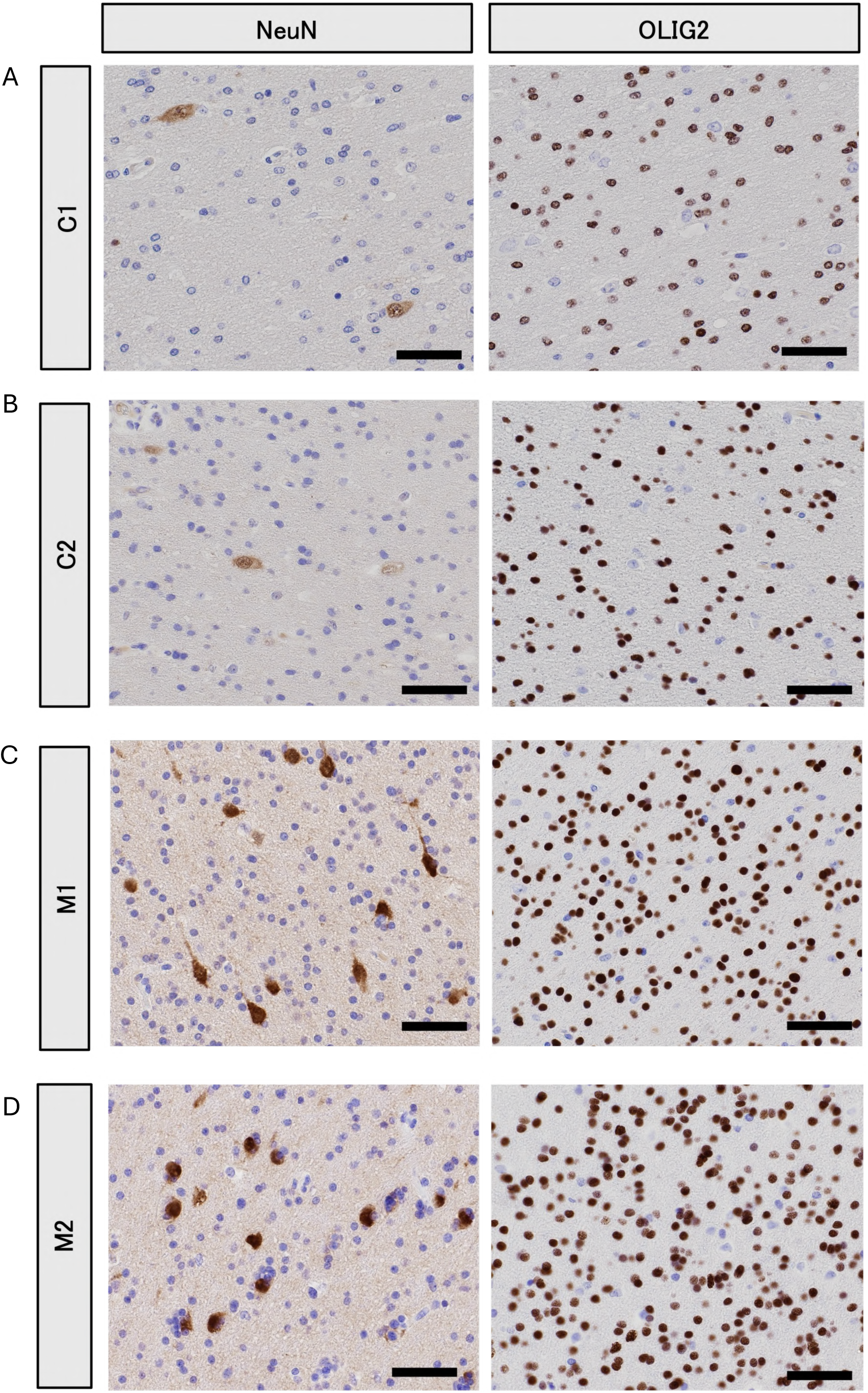
Higher-magnification histopathological images of individual control and MOGHE cases. A-D Higher-magnification images of WM regions of cortical sections from control (C1, C2) and MOGHE (M1, M2) cases, shown in A-D. Scale bars, 50 µm.

**Supplementary Fig. S2:**
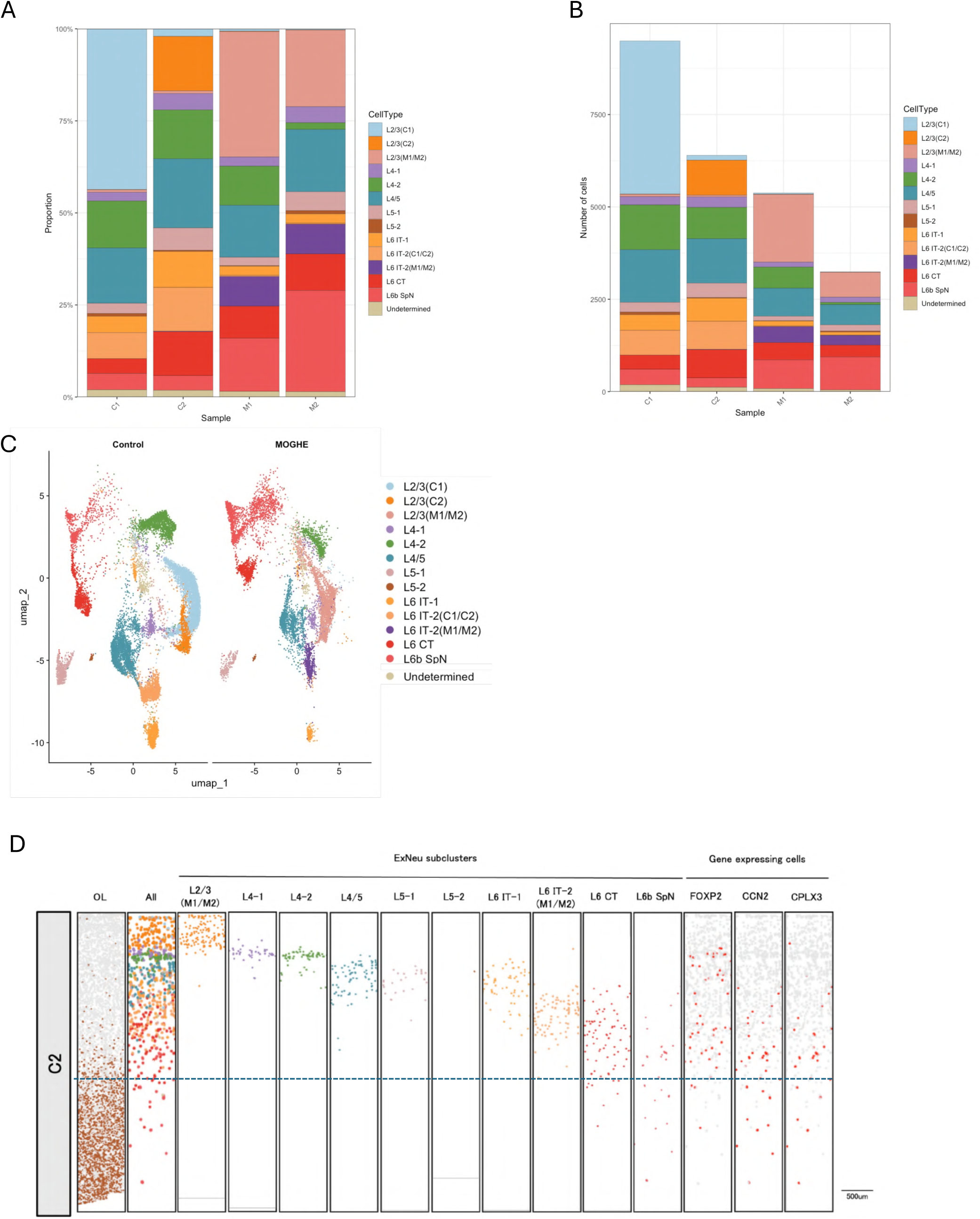
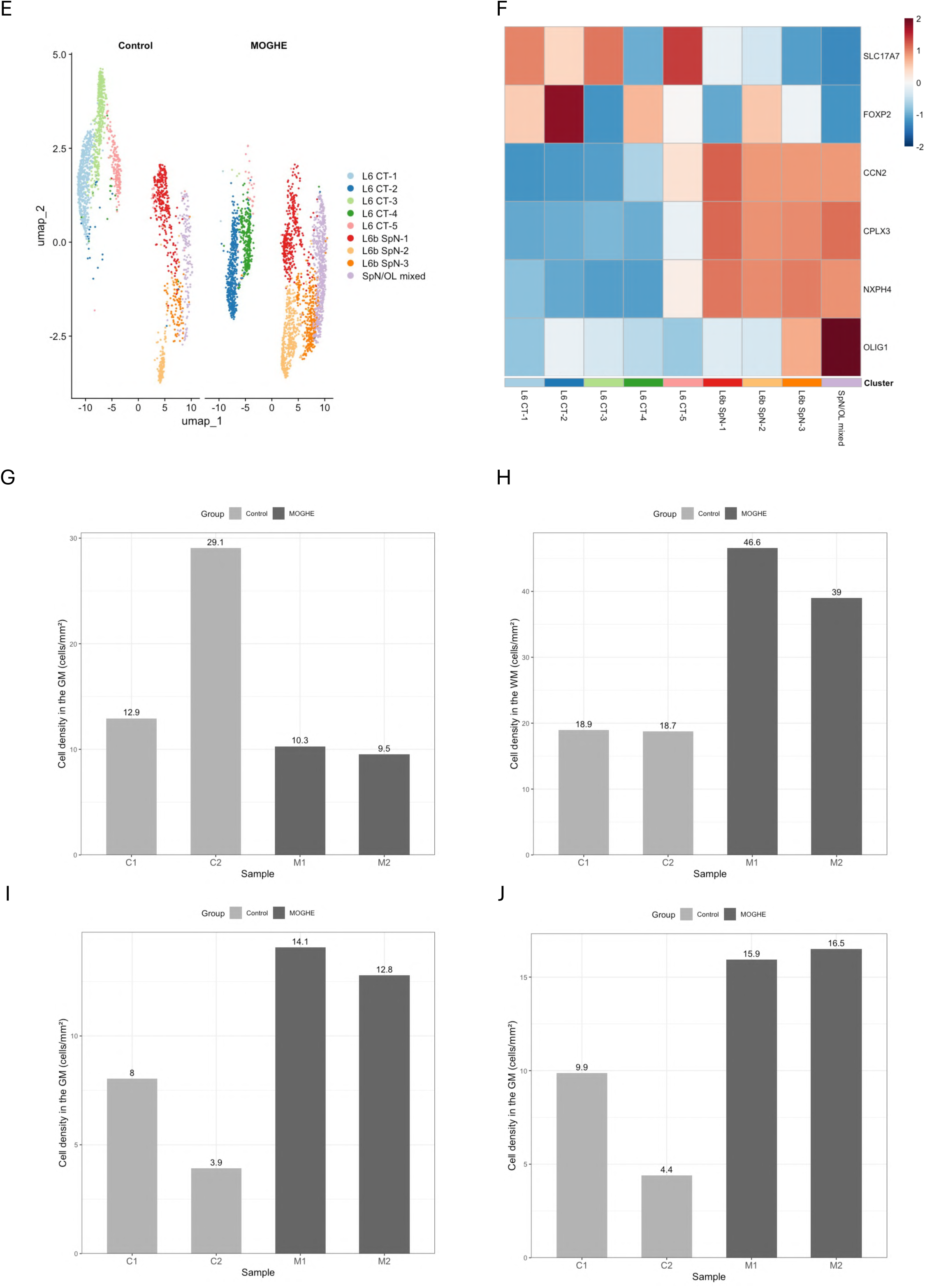
Supplementary characterization of excitatory neuron subpopulations in MOGHE. A Proportions of excitatory neuron subclusters identified by subclustering of the initial ExNeu1-7 clusters. Four glial-marker-expressing subclusters and the undetermined subcluster were excluded from this analysis (Table 3). B Absolute cell counts of excitatory neuron subclusters identified by subclustering of the initial ExNeu1-7 clusters. Four glial-marker-expressing subclusters and the undetermined subcluster were excluded from this analysis (Table 3). C UMAP visualization of excitatory neuron subclusters identified by subclustering of the initial ExNeu1-7 clusters. Four glial-marker-expressing subclusters and the undetermined subcluster were excluded from this analysis (Table 3). D Spatial maps of ExNeu subclusters in control (C2) cortical section, displayed as in Fig. 2C-E. E UMAP visualization of subclusters identified by secondary clustering of L6 CT and L6b SpN clusters. F Heatmap showing scaled expression of L6 CT- and L6b SpN-associated marker genes across subclusters identified by the secondary clustering. G Density of GM-localized L6 CT-like neurons in each sample, calculated from L6 CT subclusters identified by secondary clustering of L6 CT and L6b SpN populations (Supplementary Fig. S2E and S2F; Supplementary Table S4). H Density of WM-localized L6b SpN-like cells in each sample, calculated from L6b SpN subclusters identified by secondary clustering of L6 CT and L6b SpN populations, including the SpN/OL mixed subcluster (Supplementary Fig. S2E and S2F; Supplementary Table S4). I Density of GM-localized L6b SpN-like neurons in each sample, calculated from L6b SpN subclusters identified by secondary clustering of L6 CT and L6b SpN populations, excluding the SpN/OL mixed subcluster (Supplementary Fig. S2E and S2F; Supplementary Table S4). J Density of GM-localized L6b SpN-like cells in each sample, calculated from L6b SpN subclusters identified by secondary clustering of L6 CT and L6b SpN populations, including the SpN/OL mixed subcluster (Supplementary Fig. S2E and S2F; Supplementary Table S4).

**Supplementary Fig. S3:**
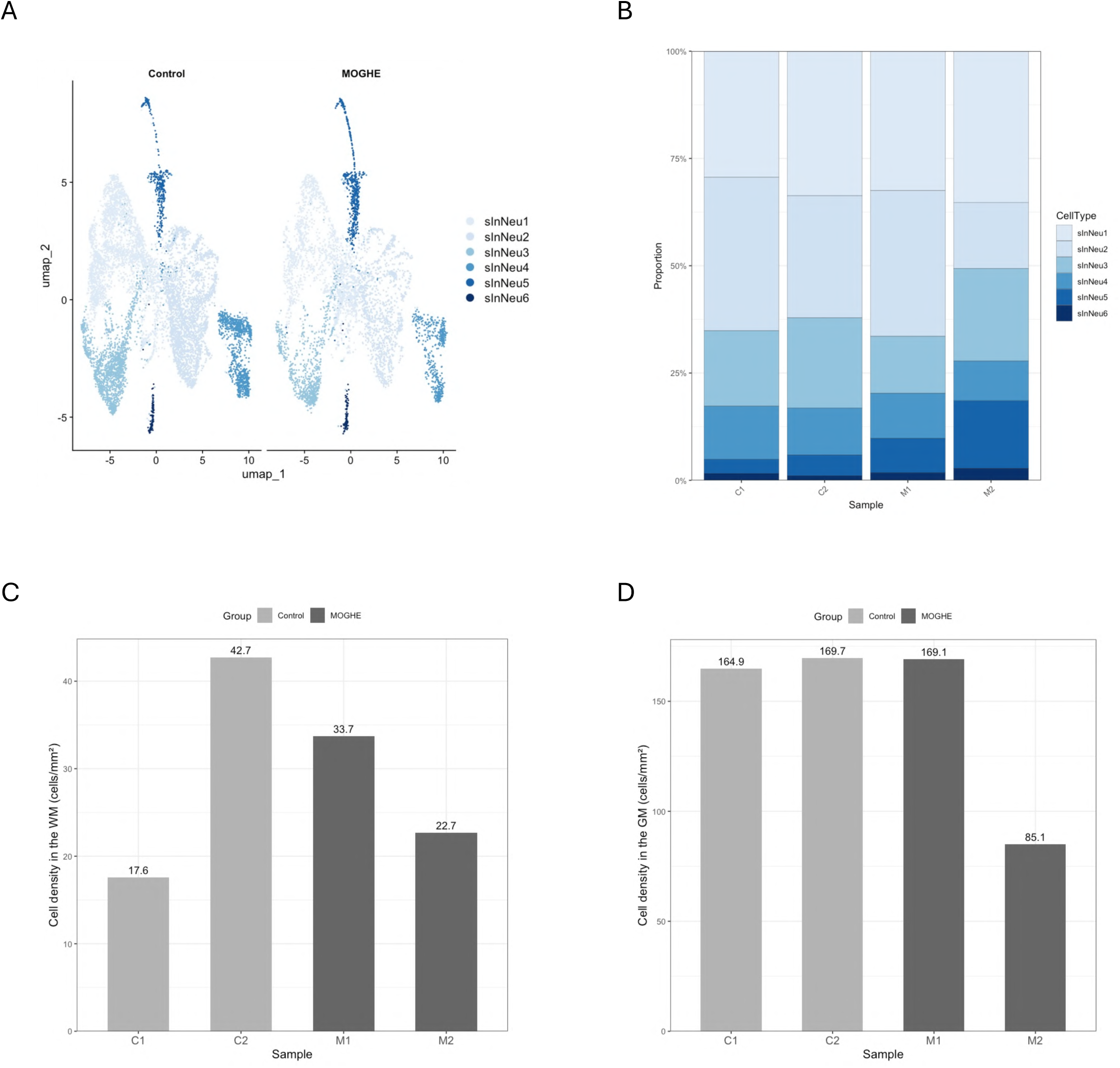
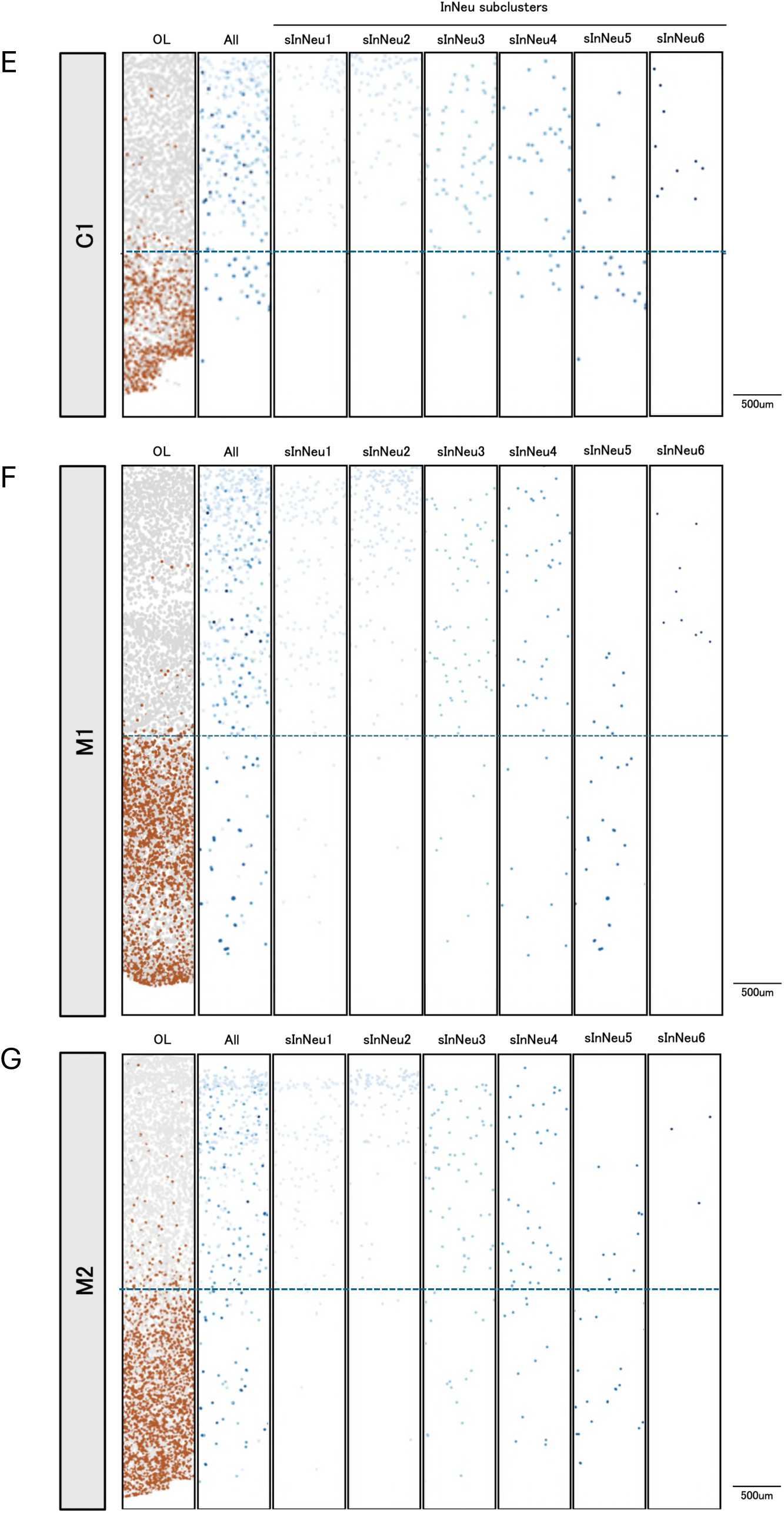
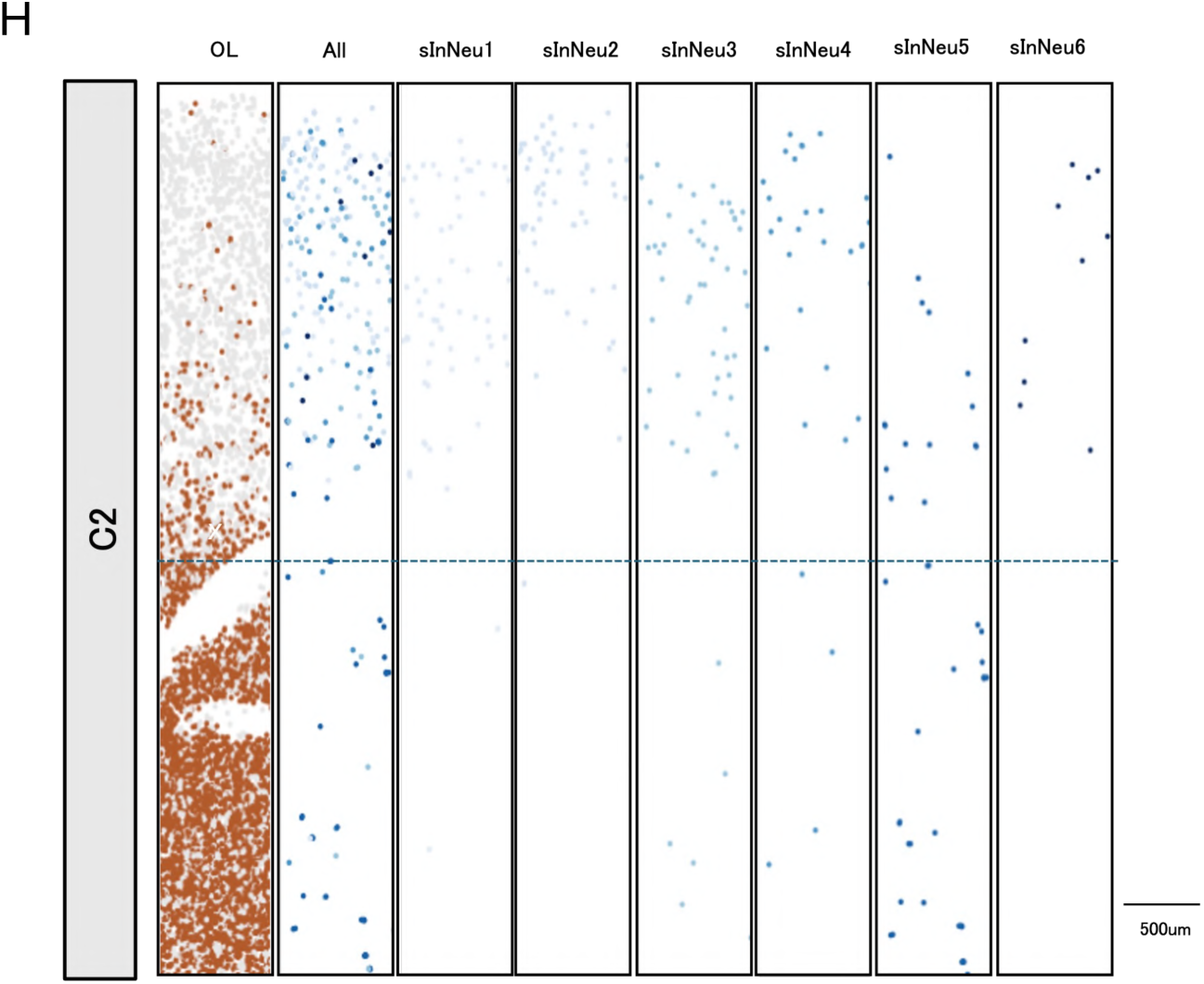
Inhibitory neuron analysis. A UMAP visualization of inhibitory neuron subclusters identified by subclustering of the initial inhibitory neuron clusters (InNeu1 and InNeu2). B Proportions of inhibitory neuron subclusters across samples. C Density of WM-localized inhibitory neurons in each sample. D Density of GM-localized inhibitory neurons in each sample. E-H Spatial maps of inhibitory neuron subclusters in control (C1, C2) and MOGHE (M1, M2) cortical sections. Dashed lines indicate the GM-WM junction. From left to right, panels show: (i) spatial distribution of OL1-OL5 clusters identified by the initial clustering, included to delineate GM and WM compartments; (ii) spatial maps of all inhibitory neuron subclusters; and (iii) spatial maps of individual InNeu subclusters.

**Supplementary Fig. S4:**
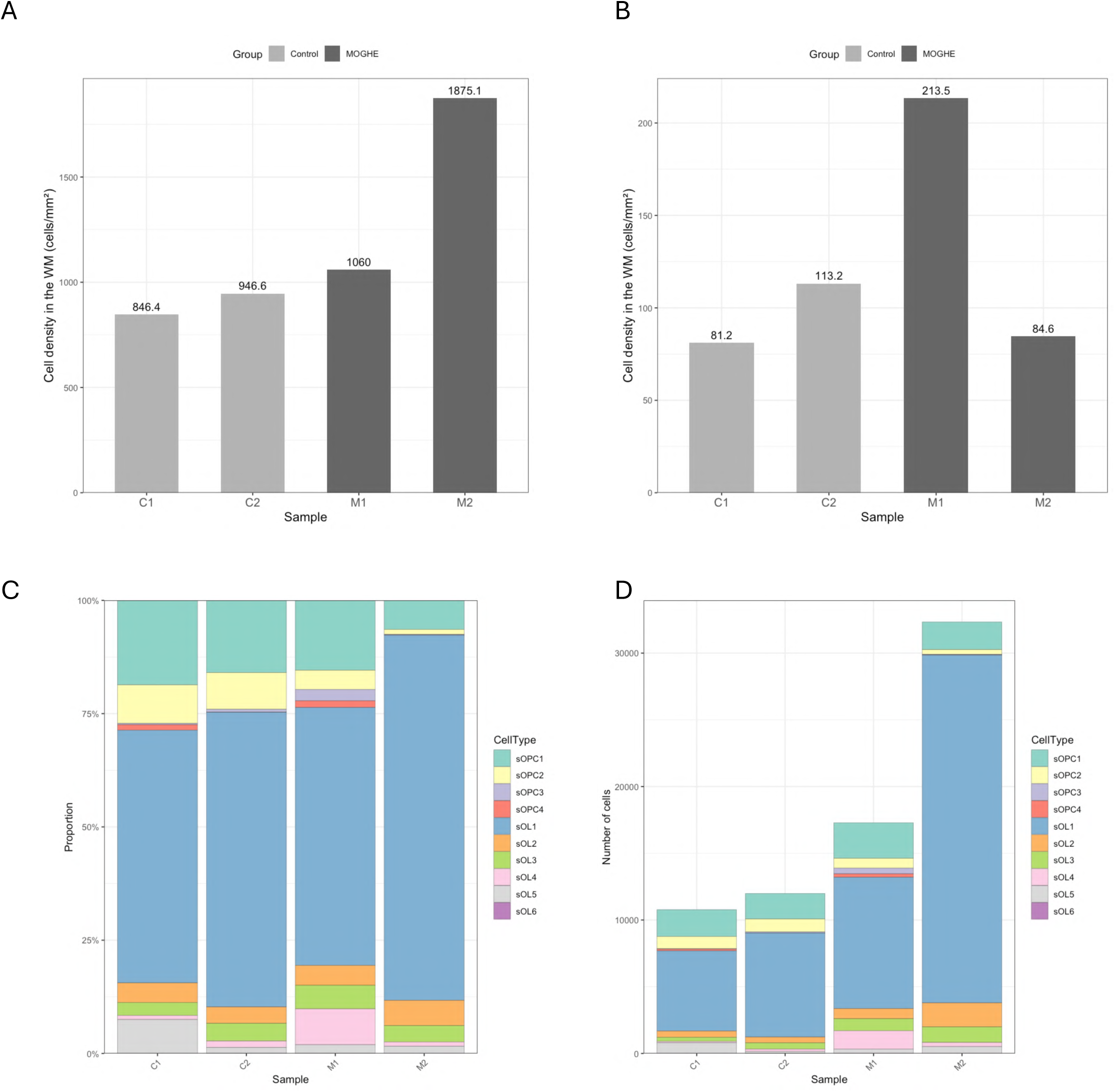
Additional characterization of oligodendroglial populations in MOGHE. A Density of WM-localized OLs in each sample. B Density of WM-localized OPCs in each sample. C Proportions of OL and OPC subclusters across samples. D Absolute cell counts of OL and OPC subclusters in each sample.

**Supplementary Fig. S5:**
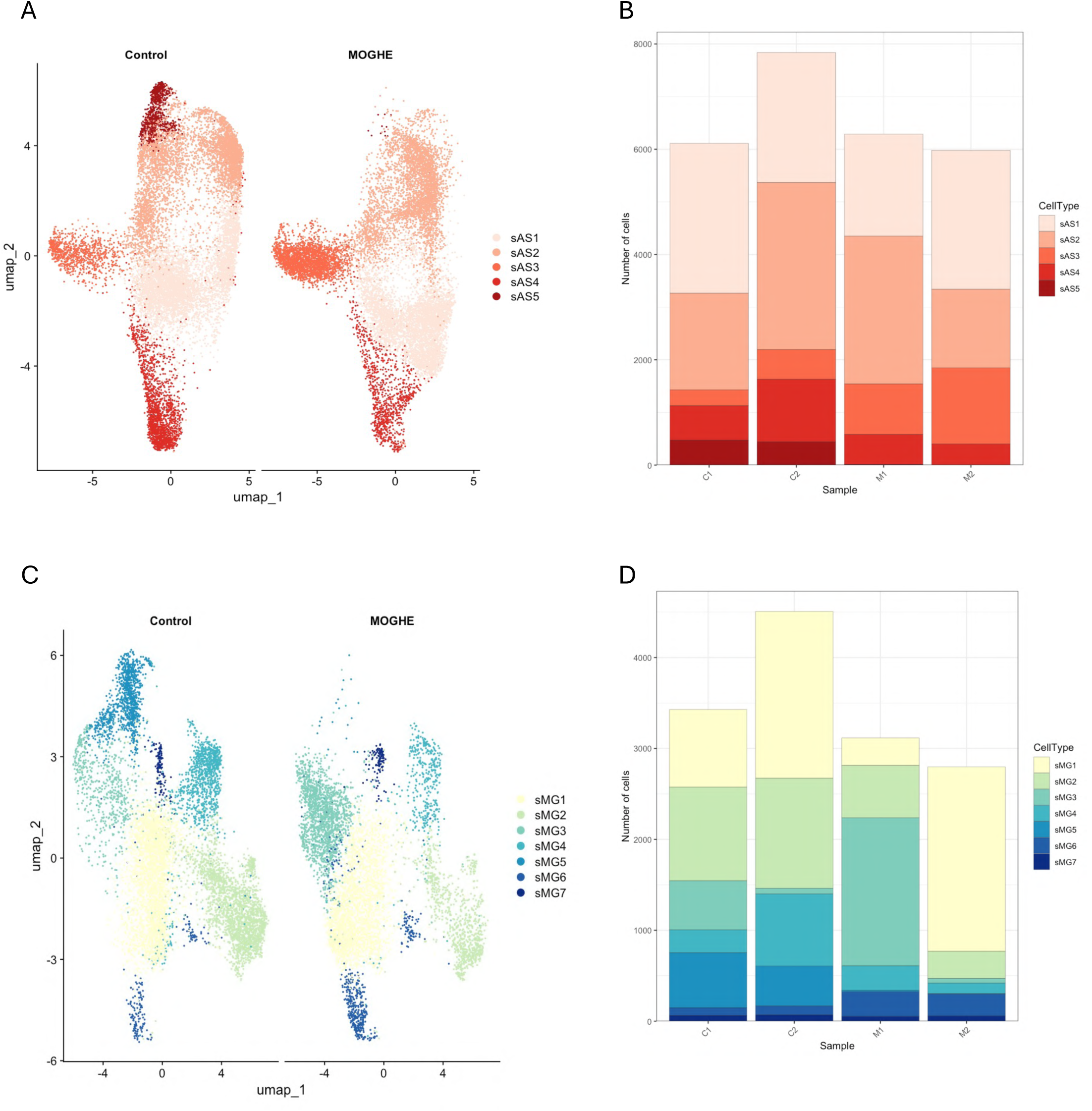
Astrocyte and microglia analyses. A UMAP visualization of astrocyte subclusters identified by the initial AS1-AS3 clusters. B Absolute cell counts of astrocyte subclusters in each sample. C UMAP visualization of microglia subclusters identified by the initial MG1-MG2 clusters. D Absolute cell counts of microglia subclusters in each sample.

**Supplementary Fig. S6:**
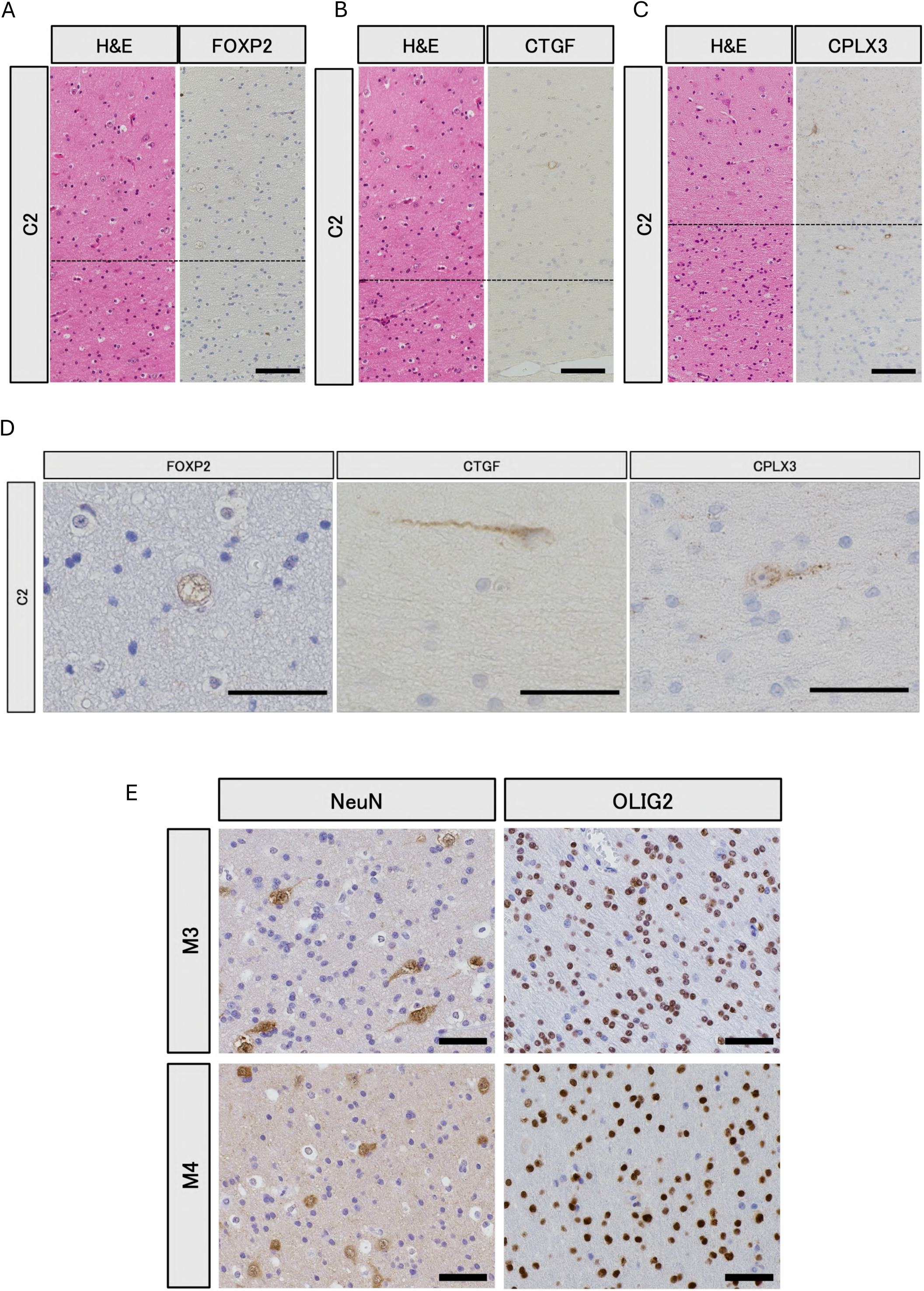
Additional histopathological and immunohistochemical analyses. A-C Immunohistochemical staining for FOXP2 (A), CTGF (B), and CPLX3 (C) in control (C2). Adjacent H&E-stained sections are shown for anatomical reference. Dashed lines indicate the GM-WM boundary. Scale bars, 100 µm. D Higher-magnification images of the WM regions shown in A-C. Scale bars, 50 µm. E Representative immunohistochemical staining for NeuN and OLIG2 in additional MOGHE cases (M3 and M4). Scale bars, 50 µm.

**Supplementary Table S1:** Quantification of NeuN-positive heterotopic neurons. NeuN-positive heterotopic neurons were counted in four representative white matter (WM) regions (Region1-Region4) for each case. Values indicate cell density (cells/mm²) for individual regions and the per-sample average.

| Sample ID | NeuN |  |  |  |  |
| --- | --- | --- | --- | --- | --- |
|  | Region1 | Region2 | Region3 | Region4 | Mean |
| C1 | 24 | 12 | 12 | 8 | 14 |
| C2 | 8 | 12 | 4 | 4 | 7 |
| M1 | 76 | 60 | 60 | 40 | 59 |
| M2 | 32 | 56 | 28 | 44 | 40 |
| M3 | 48 | 36 | 36 | 16 | 34 |
| M4 | 40 | 20 | 36 | 36 | 33 |

**Supplementary Table S2:** Quantification of OLIG2-positive cells. OLIG2-positive cells were quantified in four representative regions of interest (Region1-Region4) within the WM for each case. Values indicate cell density (cells/mm²) for individual regions and the per-sample average.

| Sample ID | OLIG2 |  |  |  |  |
| --- | --- | --- | --- | --- | --- |
|  | Region1 | Region2 | Region3 | Region4 | Mean |
| C1 | 1344 | 1656 | 1488 | 1120 | 1402 |
| C2 | 1704 | 1880 | 1572 | 1624 | 1695 |
| M1 | 3176 | 3596 | 3484 | 3000 | 3314 |
| M2 | 3060 | 3328 | 3216 | 3484 | 3272 |
| M3 | 4148 | 4132 | 3548 | 3972 | 3950 |
| M4 | 2140 | 2220 | 4460 | 2040 | 2715 |

**Supplementary Table S3:** Summary of quality control (QC) metrics. Summary of total detected cells, median genes per cell, median transcripts per cell, gray matter (GM) area and WM area across all samples.

| Sample | Total cells | Median genes per cell | Median transcripts per cell | WM area (mm<sup>2</sup>) | GM area (mm<sup>2</sup>) |
| --- | --- | --- | --- | --- | --- |
| C1 | 40612 | 207 | 252 | 7.178916 | 27.75652 |
| C2 | 41568 | 277 | 343 | 8.755295 | 19.3673 |
| M1 | 40195 | 180 | 213 | 11.869316 | 13.9996 |
| M2 | 54425 | 173 | 205 | 14.380964 | 19.93483 |

Supplementary Table S4: Cell counts for layer 6 corticothalamic (L6 CT) and layer 6b subplate neuron (L6b SpN) subclusters identified by secondary clustering.

Supplementary Table S5. Cell counts for inhibitory neuron subclusters.

Supplementary Table S6. Genes enriched in MOGHE oligodendroglial subclusters compared with Control oligodendroglial subclusters (p_adj < 0.05, log2FC > 0). (Excel)

Supplementary Table S7. Cell counts for astrocyte subclusters.

Supplementary Table S8. Cell counts for microglia subclusters.

Supplementary Table S9. Genes enriched in WM-localized excitatory neuron (ExNeu) subclusters compared with GM-localized ExNeu subclusters (p_adj < 0.05, log2FC > 0)

Supplementary Table S10. Genes decreased in WM-localized ExNeu subclusters compared with GM-localized ExNeu subclusters (p_adj < 0.05, log2FC < 0)

Supplementary Table S11. Genes enriched in WM-localized L6 CT and L6b SpN subclusters compared with GM-localized L6 CT and L6b SpN subclusters in MOGHE samples (p_adj < 0.05, log2FC > 0). (Excel)

Supplementary Table S12. Genes decreased in WM-localized L6 CT and L6b SpN subclusters compared with GM-localized L6 CT and L6b SpN subclusters in MOGHE samples (p_adj < 0.05, log2FC < 0) (Excel)

**Supplementary Table S13.** Antibodies used in this study.

| Target | Host | Manufacturer | cat | lot | Human IHC | Mouse IHC |
| --- | --- | --- | --- | --- | --- | --- |
| NeuN | Mouse | Millipore | #MAB377 | – | ○ | – |
| OLIG2 | Rabbit | IBL | #18953 | – | ○ | – |
| NeuN | Guinea Pig | Millipore | #ABN90 | #4208775 | – | ○ |
| TLE4 | Mouse | SantaCruz | #sc-365406 | #G0324 | – | ○ |
| FOXP2 | Rabbit | Abcam | #ab16046 | #1133400-4 | ○ | – |
| CPLX3 | Rabbit | Abcam | #ab308463 | #1047790-7 | ○ | – |
| CPLX3 | Rabbit | Synaptic Systems | #122302 | #2-12 | – | ○ |
| CTGF | Mouse | SantaCruz | #sc-365970 | #C2724 | ○ | – |

## Methods

### Human brain samples and ethics

Human brain tissue samples were obtained from patients with medically refractory epilepsy who underwent surgical resection at the National Center Hospital, National Center of Neurology and Psychiatry, Kodaira, Tokyo, Japan (NCNP). Two patients were histopathologically diagnosed with MOGHE based on established quantitative criteria, including increased densities of OLIG2-positive cells (>2,200 cells/mm²) and NeuN-positive neurons in the white matter (>30 cells/mm²), as previously described.^22^ Cell counts were performed using automated cell detection in QuPath (v0.5.1) and confirmed by experienced neuropathologists.^39^ As controls, cortical tissue was obtained from age- and sex-matched patients with mTLE associated with HS, using histopathologically normal cortical regions.

All clinical information and tissue specimens were collected in accordance with the principles of the Declaration of Helsinki. Written informed consent was obtained from all patients or their legal guardians prior to surgery. This study was reviewed and approved by the Ethics Committee of the National Center of Neurology and Psychiatry, Japan (NCNP-A2018-050).

For iST experiments, surgically resected brain tissues were fixed in 4% paraformaldehyde for 48 hours, followed by dehydration through a graded ethanol series, clearing in xylene, and embedding in paraffin. Formalin-fixed paraffin-embedded (FFPE) sections were subsequently used for downstream analyses.

### Genome analysis

Targeted genomic analysis was performed to detect somatic variants in *SLC35A2* and other genes implicated in malformations of cortical development (MCD), including components of the PI3K-AKT-mTOR signaling pathway.

Genomic DNA (20 ng) extracted from frozen brain tissue was subjected to multiplex PCR amplification using the AmpliSeq HD Library Kit (Thermo Fisher Scientific, USA), according to the manufacturer’s instructions. Barcoded libraries were quantified, pooled at equimolar concentrations, and processed on an Ion Chef Instrument (Thermo Fisher Scientific) for template preparation. Libraries were then loaded onto Ion 540 chips (four samples per chip) and sequenced on an Ion GeneStudio S5 system (Thermo Fisher Scientific) with a read length of 200 bp and 500 flow cycles.

### Xenium spatial transcriptomics

Imaging-based spatial transcriptomic analysis was performed using the Xenium In Situ Gene Expression platform (10x Genomics). FFPE brain tissues were sectioned at a thickness of 5 µm and mounted onto Xenium slides. Sections were deparaffinized and decrosslinked following the manufacturer’s protocols.

Probe hybridization, ligation, and signal amplification were carried out according to the Xenium In Situ Gene Expression user guidelines (10x Genomics). Spatial transcript profiling was conducted using the Xenium Prime 5K Human Pan Tissue and Pathways Panel, which targets 5,001 genes. Following probe detection, cell segmentation staining was performed using the Xenium Cell Segmentation Staining Reagents Kit, with incubation carried out overnight at 4 ° C. Autofluorescence quenching and nuclear staining were subsequently performed under light-protected conditions.

Slides were imaged using the Xenium Analyzer (10x Genomics) to acquire high-resolution spatial gene expression data. After completion of Xenium imaging, probe signals were removed using a quencher removal solution, and sections were washed according to the manufacturer’s demonstrated protocol. Tissue sections were then counterstained with H&E to visualize cortical architecture. Whole-slide images were acquired using a NanoZoomer S60 digital slide scanner (Hamamatsu Photonics, Japan).

### Xenium data analysis

Cell segmentation was performed using the default nuclear- and cell-boundary-based segmentation pipeline implemented in the Xenium Analyzer (10x Genomics). Downstream analyses were conducted using the Seurat package (v5.2.1) in R.^40^

To remove low-quality segments, cells were filtered based on the number of detected transcripts and genes, retaining cells with nFeature_Xenium > 50 and nCount_Xenium > 100. Filtered data were normalized using the NormalizeData function in Seurat. Dimensionality reduction was performed by principal component analysis (PCA), and the resulting principal components were used to construct a shared nearest neighbor (SNN) graph with FindNeighbors. Unsupervised clustering was performed using FindClusters, and clusters were visualized by uniform manifold approximation and projection (UMAP) using RunUMAP.

Major brain cell types, including excitatory neurons, inhibitory neurons, oligodendrocytes, oligodendrocyte precursor cells, astrocytes, microglia, and endothelial cells, were annotated based on the expression of established marker genes together with spatial distribution patterns. To resolve finer transcriptional heterogeneity, major cell types̶including excitatory neurons, inhibitory neurons, oligodendrocytes, oligodendrocyte precursor cells, microglia and astrocytes̶were further subsetted and independently reclustered using cell type-specific principal components and clustering parameters. Differentially expressed genes were identified using the FindMarkers function in Seurat.

Spatial visualization and alignment of transcriptomic data with histological images were performed using Xenium Explorer (v4.1.0), based on nuclear staining images. H&E-stained images were overlaid with spatial transcriptomic data to enable integrated interpretation of molecular and histological features.

### Immunohistochemistry (human brain tissue)

FFPE human brain samples were sectioned at a thickness of 6 µm. Immunohistochemistry (IHC) staining was performed using standard protocols with minor modifications. Briefly, sections were deparaffinized and endogenous peroxidase activity was quenched by incubation in methanol containing hydrogen peroxide. Antigen retrieval was performed by heat-induced epitope retrieval in sodium citrate buffer. After blocking with normal serum, sections were incubated with primary antibodies overnight at 4 ° C. Immunoreactivity was detected using a polymer-based detection system and visualized with diaminobenzidine. Sections were counterstained with hematoxylin, dehydrated, and mounted for microscopic analysis. Primary antibodies used in this study are listed in Supplementary Table S13.

### Animals

All animal experiments were approved by the Animal Care and Use Committee of the National Center of Neurology and Psychiatry, Japan, and were conducted in accordance with the NIH Guide for the Care and Use of Laboratory Animals. Mice were bred and maintained in a temperature- and humidity-controlled, specific pathogen-free facility under a 12-hour light/dark cycle, with ad libitum access to food and water.

Conditional *Slc35a2* knockout mice were generated using a floxed *Slc35a2* allele. The *Slc35a2* floxed strain (C57BL/6JGpt-Slc35a2em1Cflox/Gpt; Strain No. T062903) was obtained from GemPharmatech (USA) and maintained on a C57BL/6J background. LoxP sites flank exon 3 of the *Slc35a2* gene, enabling Cre-mediated conditional deletion.

For conditional gene deletion in dorsal pallium-derived lineages, *Slc35a2*-floxed mice were crossed with Emx1-Cre driver mice (RIKEN BRC, Japan; Stock No. RBRC00808).^26^ Female *Slc35a2*^fl/+^; Emx1-Cre-positive mice were designated as cKO mice, and *Slc35a2*^fl/+^; Cre-negative littermates were used as controls.

### Immunohistochemistry (mouse brain tissue)

Two-month-old control and cKO female mice were deeply anesthetized and perfused transcardially with phosphate-buffered saline (PBS, 0.1 M, pH 7.4), followed by 4% paraformaldehyde (PFA) in PBS. Brains were collected and post-fixed overnight in the same fixative. Tissues were then cryoprotected by sequential immersion in 10%, 20%, and 30% sucrose in PBS, embedded in optimal cutting temperature (OCT) compound (Cat#4583, Sakura Finetek, Japan), and sectioned coronally at 20 µm thickness using a cryostat (Leica, Germany).

For immunohistochemistry, sections were first bleached in 0.1% H_2_O_2_ in PBS at room temperature to quench endogenous peroxidase activity. Following antigen retrieval when required (Dako, Cat#S1699, Japan), the sections were permeabilized and blocked in 0.2% Tween-20 in PBS (PBST) containing 4% normal donkey serum and 1% bovine serum albumin, and then incubated with primary antibodies overnight at 4 °C. After washing, sections were incubated with biotinylated secondary antibodies at room temperature, followed by incubation with VECTASTAIN ABC reagent according to the manufacturer’s instructions (ABC Standard Kit, Cat#PK-4000, Vector Laboratories, USA). Immunoreactivity was visualized using 0.05% 3,3’-diaminobenzidine (DAB) with 0.003% H_2_O_2_ in PBS. Sections were then mounted for microscopic analysis. Primary antibodies used in this study are listed in Supplementary Table S13.

### Quantitative analysis of heterotopic neurons in mice

Images of immunostained coronal sections were acquired using a Keyence microscope system (Keyence, USA). To quantify cell density within the WM, the area of cg was defined according to the coronal atlas in the Allen Mouse Brain Atlas 3-D Viewer.^41^ NeuN-, TLE4-, and CPLX3-positive cells within the cg were manually counted using Fiji software (v2.16.0).^42^ Cell density was calculated as the number of positive cells per unit area of the cg. Data are presented as mean ± SEM. Statistical comparisons between control and cKO mice were performed using Welch’s t-test in Microsoft Excel (Microsoft Corporation, USA). P values < 0.05 were considered statistically significant.

## Data availability

Raw sequence file and spatial transcriptomic data obtained in this study will be available after publication. Before that, all data will be shared upon reasonable requests.

## Code availability

Proccessing worlfows used for all analyses are available on Github at https://github.com/Hoshino-lab/Tabe_2026

## Acknowledgments

We thank Drs. Takahiro Hayashi and Yuiko Kimura (National Institute of Neurology and Psychiatry, Japan) for their contribution to the collection of human surgical specimens, Yuji Nakayama, Kazumasa Sekiguchi, and Takahito Ichikawa (National Institute of Neurology and Psychiatry, Japan) for their contribution to pathological diagnosis, and Chiaki Ohtaka-Maruyama (Tokyo Metropolitan Institute of Medical Science, Japan) for fruitful discussions.

## Author Contributions

N.N.K.T., S.M., K.Y., M.I., and M.H. conceived and designed the study. K.I. and M.I. coordinated patient recruitment and neurosurgical tissue collection. N.N.K.T., S.M., K.S., and K.N. contributed to transcriptomic data generation and analysis. K.Y., E.U., T.S., and M.T. contributed to tissue processing. K.Y. performed human histological and immunohistochemical analyses. N.N.K.T., A.H., and S.O. contributed to mouse breeding and husbandry. N.N.K.T. performed mouse histological and immunohistochemical analyses. K.I., K.M., and K.K. performed genome analysis. N.N.K.T. and S.M. wrote the original draft. M.S., M.I., and M.H. reviewed and edited the manuscript. All authors reviewed and approved the final manuscript.

